# Alcohol Cues Elicit Different Abnormalities in Brain Networks of Abstinent Men and Women with Alcohol Use Disorders

**DOI:** 10.1101/2021.03.31.437778

**Authors:** Kayle S. Sawyer, Marlene Oscar-Berman, Susan Mosher Ruiz, Ksenija Marinkovic, Mary M. Valmas, Gordon J. Harris

## Abstract

We employed fMRI in 84 men and women with and without a history of alcohol use disorders (ALC and NC, respectively), to explore how gender interacts with alcoholism as reflected in brain activity elicited by alcohol cues. Brain activation was measured in a working memory task (delayed matching-to-sample) with emotional faces as the sample and match cues. During the delay period, intervening distractors were either reward-salient cues (alcoholic beverages) or neutral cues (nonalcoholic beverages or scrambled pictures). ALC women (ALCw) had higher accuracy than ALC men (ALCm). Analyses of scans during the viewing of distractor images revealed significant group-by-gender interactions. Compared to NC men, ALCm evidenced lower activation contrast between reward-salient cues and neutral cues in *default mode network* regions (including superior prefrontal and precuneus areas), while ALCw had more activation than NC women. Similar interactions were observed for *task-regions* (including superior parietal, lateral occipital, and prefrontal areas). Region of interest analyses showed that the ALC group had significantly higher levels of activation throughout reward-related circuitry during alcohol distractor interference than during scrambled picture interference. These results suggest that abstinent ALCm and ALCw differ in processing reward-salient cues, which can impact treatment and recovery.

**Highlights:** Brain reward regions activate highly when individuals with a history of alcohol use disorders view alcoholic beverages.

The brain regions identified subserve vision, memory, and judgement.

Opposite abnormalities in activation patterns appeared for alcoholic men and women.

## 1. Introduction

Alcohol use disorders (AUD) have been associated with deficits in cognitive and emotional functions (Oscar-Berman *et al*., 2014). Because of their reward salience, alcohol cues such as pictures of alcoholic beverages elicit attentional bias and brain activation in individuals with AUD (Carter & Tiffany, 1999; Goldstein & Volkow, 2002; Schacht *et al*., 2013). Alcohol cues induce a hyperattentive state with attention drawn to the rewarding stimuli (Townshend & Duka, 2001; Franken, 2003; Field & Cox, 2008; Vollstädt-Klein *et al*., 2011). Therefore, the cues selectively interfere with other cognitive abilities such as memory. Importantly, attentional bias toward alcohol-related stimuli also has been associated with level of craving, consumption, dependence, and physiological arousal (Sharma *et al*., 2001; Ryan, 2002; Field *et al*., 2004; Field & Eastwood, 2005; Bordnick *et al*., 2008; Sinha *et al*., 2009; Wiers *et al*., 2014; Sawyer *et al*., 2015).

Functional magnetic resonance imaging (fMRI) studies of attentional biasing, and specifically cue sensitivity, have often included either only men with and without a history of AUD (ALC and NC groups), or groups of men and women with sample sizes too small to examine gender effects (Fryer *et al*., 2013; Schacht *et al*., 2013). However, gender impacts the ways in which alcohol affects the brain and behavior (Becker *et al*., 2017; Sawyer *et al*., 2017, 2018, 2019; Seitz *et al*., 2017; Rivas-Grajales *et al*., 2018; Hoffman *et al*., 2019; Kaag *et al*., 2019; Fama *et al*., 2020; Verplaetse *et al*., 2021), due to interactions with physiological and social factors (Ruiz & Oscar-Berman, 2015; Mosher Ruiz *et al*., 2017). In the present study, we examined gender differences using a delayed matching-to-sample (DMTS) task (Dolcos & McCarthy, 2006) with alcohol cues serving as distractor stimuli, in a cohort of ALC and NC men and women (AUDm, AUDw, NCm, and NCw). The DMTS task requires an attention-demanding kind of memory called “working memory.” In this study, the participants were required to remember photographs of emotional faces while distracting pictures of alcoholic beverages, nonalcoholic beverages, or scrambled images intervened during the delay period. We chose faces as the sample stimuli for two primary reasons. First, in a previous study (Marinkovic *et al*., 2009) we found that men with AUD had abnormally low brain activity in temporal limbic regions when viewing faces, and second, we used the same dataset that we had acquired for a prior report (Oscar-Berman *et al*., 2019) in which we described the brain’s responses to the initial to-be-encoded emotional faces phase (the sample) of the DMTS task. For the present study, the data derived from the delay and match portions of the task allowed us to test the attentional biasing effect, wherein we expected alcohol cues, more than other cue types, to impair performance on memory for face identity.

Functional MRI tasks activate multiple brain networks, and abnormalities in the default mode network (DMN) have been implicated in AUD and in psychiatric disorders (Menon, 2011; Zhang & Volkow, 2019). The DMN has been observed to be more active during story telling, reading and memory tasks, imagining future scenarios, self-reference, rumination, and when the mind wanders while staring at a fixation cross during fMRI scanning (Tops *et al*., 2014; Beaty *et al*., 2016; Buckner & DiNicola, 2019). We refer to the DMN regions as *fixation-regions* because they are more active during the idle delay intervals when the fixation stimulus is presented between DMTS trials than during stimulus presentations. Vertex-wise analyses have revealed that cortical fixation-regions include: (1) an “anterior hub,” consisting of portions of the rostral anterior cingulate cortex (ACC), ventromedial prefrontal cortex, and medial superior frontal cortex; (2) a “posterior hub,” which includes portions of the posterior cingulate and precuneus, (3) the temporoparietal junction, which covers parts of the angular gyrus and inferior parietal lobule; and (4) the superior and middle temporal gyrus region (Margulies *et al*., 2016; Buckner & DiNicola, 2019; Uddin *et al*., 2019).

In addition to the DMN regions, we used vertex-wise analyses to examine *task-regions*, which are more engaged during the DMTS task than while looking at unengaging fixation crosses. Literature on distractor interference during working memory has suggested that task-regions involve a distributed network including prefrontal cortex, along with dorsal and ventral visual association cortex, which are necessary for attentional functioning (Loeber *et al*., 2009) and for inhibiting distracting visual stimuli (Jha *et al*., 2004; Clapp *et al*., 2010). Additional task-regions involved in attention, working memory, and emotional processing, include the dorsal ACC and lateral prefrontal areas. The dorsal ACC in particular has been implicated in craving and attentional biasing (Goldstein & Volkow, 2011).

In advance of any analyses, we used prior literature to select ten *a priori* anatomically-defined regions of interest (ROI) involved in alcohol cue exposure, distractor interference, craving, reward processing and salience, or working memory for emotional faces. The first nine ROI are the dorsolateral prefrontal cortex (DLPFC), ventrolateral prefrontal cortex (VLPFC), orbitofrontal cortex (OFC), insular cortex, parahippocampal gyrus, hippocampus, amygdala, fusiform, and ACC. Previous studies provide support for each of those nine a priori ROI (George *et al*., 2001; Wrase *et al*., 2002; Tapert *et al*., 2003; Myrick *et al*., 2004; Heinz *et al*., 2007; Ray *et al*., 2010; Goldstein & Volkow, 2011; Schacht *et al*., 2013; Field *et al*., 2014; Alba-Ferrara *et al*., 2016; Sawyer *et al*., 2020). The tenth ROI is the “extended reward and oversight system” (EROS) as described and named in our previous papers (Makris *et al*., 2008; Sawyer *et al*., 2017). The EROS ROI is a single large but discontinuous composite ROI that had been created by combining 11 regions (seven of the ROI noted above, all but the VLPFC and fusiform), plus an additional four: nucleus accumbens, ventral diencephalon, subcallosal cortex, and temporal pole. For each of the ten ROI, we intended to confirm findings from each of the aforementioned studies that had identified abnormal activation by alcohol cues in AUD, and additionally to investigate differences between men and women. We expected lower brain activation in the ALC group in regions involved in facial identity and inhibition of distractor interference, but higher in regions responsible for reward salience.

In summary, we investigated brain activation for ten ROI, and for vertex-wise cortical analyses of fixation-regions and task-regions. We examined the accuracy of the participants’ memory for the face identities after exposure to attentionally salient pictures to test our hypothesis that alcohol cues would distract the ALC group more than the NC group. We determined how brain regions were activated by the distractor contrasts, how the contrasts differed for ALC and NC groups, and how those abnormalities varied by gender. We hypothesized that attentional biasing would be evident for the ALC group in the form of stronger brain activity contrasts (alcoholic beverage cues compared to nonalcoholic and scrambled stimuli). Regarding gender differences, we made predictions based upon previous work in our laboratory wherein we found that brain regions of ALCw (compared to NCw) were overactive in response to highly charged emotional stimuli (Sawyer *et al*., 2019). We hypothesized that the ALCw would evidence hyperactivation to emotionally valent stimuli, whereas the activation contrasts for ALCm would be weaker. We also expected to replicate prior results (Marinkovic *et al*., 2009; Sawyer *et al*., 2019) showing lower responses in ALCm than NCm.

## 2. Materials and Methods

### 2.1 Participants

Participants in this study included 42 abstinent long-term ALC individuals (21 ALCw) and 42 NC controls (21 NCw), with comparable age, education, and IQ (see Table 1 in the Results). Participants were recruited through flyers placed in treatment and after-care facilities, the Boston VA Healthcare System facility, Massachusetts General Hospital, the Boston University School of Medicine, and in public places (*e.g.*, churches, stores), as well as through newspaper and internet advertisements. This study was reviewed and approved by human studies Institutional Review Boards at the affiliated institutions. All participants gave written informed consent prior to participation, and they were compensated for their time.

**Table 1.** Participant characteristics. Participants Characteristics (*p* < 0.05): ^a^Control Women > Control Men; ^b^Control Men > Alcoholic Men; ^c^Alcoholics > Controls; ^d^Alcoholic Men > Control Men; ^e^Alcoholic Women > Control Women; ^f^Alcoholic Men > Alcoholic Women; ^g^Alcoholic Women > Alcoholic Men. See Results for additional details on number of participants. Abbreviations: WAIS = Wechsler Adult Intelligence Scale; WMS = Wechsler Memory Scale; IMI = Immediate Memory Index; DMI = Delayed Memory Index. Five NCm and two NCw reported being lifetime abstainers, and one NCw was unable to report an accurate length of sobriety.

|  | ALCOHOLICS |  |  | CONTROLS |  |  |
| --- | --- | --- | --- | --- | --- | --- |
|  | ALC | ALCw | ALCm | NC | NCw | NCm |
|  | *n* = 42 | *n* = 21 | *n* = 21 | *n* = 42 | *n* = 21 | *n* = 21 |
| **Age<sup>a</sup> (years)** |  |  |  |  |  |  |
| *mean* | 53.9 | 53.4 | 54.4 | 53.9 | 57.7 | 50.2 |
| *standard deviation* | 11.0 | 11.4 | 10.8 | 12.4 | 13.6 | 10.1 |
| *range* | 26.5 - 76.7 | 26.5 - 73.0 | 26.6 - 76.7 | 25.8 - 76.9 | 25.8 - 76.9 | 29.0 - 69.6 |
| **Education<sup>b</sup> (years)** |  |  |  |  |  |  |
| *mean* | 14.7 | 15.3 | 14.1 | 15.5 | 15.6 | 15.4 |
| *standard deviation* | 2.0 | 2.0 | 1.9 | 2.0 | 2.3 | 1.6 |
| *range* | 12 - 19 | 12 - 19 | 12 - 18 | 12 - 19 | 12 - 20 | 12 - 18 |
| **WAIS-III Full Scale IQ** |  |  |  |  |  |  |
| *mean* | 110.3 | 110.1 | 110.5 | 111.6 | 111.2 | 112.0 |
| *standard deviation* | 15.0 | 14.2 | 16.0 | 16.3 | 19.3 | 13.1 |
| *range* | 72 - 140 | 72 - 137 | 81 - 140 | 79 - 152 | 79 - 142 | 90 - 152 |
| **WMS-III IMI** |  |  |  |  |  |  |
| *mean* | 109.7 | 114.4 | 104.7 | 111.9 | 114.8 | 109.0 |
| *standard deviation* | 16.6 | 18.3 | 13.4 | 16.9 | 16.4 | 17.4 |
| *range* | 63 - 144 | 63 - 144 | 82 - 130 | 80 - 146 | 84 - 138 | 80 - 146 |
| **WMS-III DMI** |  |  |  |  |  |  |
| *mean* | 112.6 | 116.7 | 108.3 | 111.8 | 113.5 | 110.1 |
| *standard deviation* | 17.3 | 20.4 | 12.5 | 16.0 | 14.9 | 17.2 |
| *range* | 52 - 140 | 52 - 140 | 86 - 132 | 83 - 150 | 83 - 140 | 84 - 150 |
| **Duration of Heavy Drinking<sup>cdef</sup> (years)** |  |  |  |  |  |  |
| *mean* | 17.4 | 14.3 | 20.5 | 0.0 | 0.0 | 0.0 |
| *standard deviation* | 7.7 | 5.2 | 8.5 | 0.0 | 0.0 | 0.0 |
| *range* | 5.0 - 35.0 | 6.0 - 25.0 | 5.0 - 35.0 | 0.0 - 0.0 | 0.0 - 0.0 | 0.0 - 0.0 |
| **Quantity Frequency Index<sup>cde</sup> (ounces ethanol/day; ~drinks/day)** |  |  |  |  |  |  |
| *mean* | 11.2 | 8.7 | 13.7 | 0.3 | 0.2 | 0.4 |
| *standard deviation* | 8.8 | 5.8 | 10.5 | 0.6 | 0.5 | 0.7 |
| *range* | 2.7 - 38.4 | 2.7 - 28.1 | 4.5 - 38.4 | 0.0 - 2.6 | 0.0 - 2.4 | 0.0 - 2.6 |
| **Length of Sobriety<sup>cde</sup> (years)** |  |  |  |  |  |  |
| *mean* | 8.3 | 10.6 | 5.9 | 2.1 | 3.6 | 0.5 |
| *standard deviation* | 10.3 | 11.1 | 8.8 | 6.4 | 8.5 | 1.3 |
| *range* | 0.1 - 32.3 | 0.1 - 32.1 | 0.1 - 32.3 | 0.002 - 29.2 | 0.002 - 29.2 | 0.002 - 5.1 |
| **Penn Alcohol Craving Scale<sup>cde</sup>** |  |  |  |  |  |  |
| *mean* | 3.8 | 3.8 | 3.9 | 1.2 | 1.1 | 1.3 |
| *standard deviation* | 4.4 | 5.1 | 3.8 | 2.0 | 2.2 | 1.8 |
| *range* | 0 - 19 | 0 - 19 | 0 - 12 | 0 - 9 | 0 - 9 | 0 - 5 |
| **Hamilton Rating Scale for Depression<sup>g</sup>** |  |  |  |  |  |  |
| *mean* | 3.5 | 4.9 | 2.2 | 2.4 | 3.1 | 1.8 |
| *standard deviation* | 4.2 | 4.1 | 4.0 | 2.8 | 3.3 | 2.1 |
| *range* | 0 - 18 | 0 - 17 | 0 - 18 | 0 - 12 | 0 - 12 | 0 - 8 |

Selection procedures included a telephone interview to determine age, education, health and alcohol and drug use history, including prescription drugs. Participants were right-handed, had normal or corrected-to-normal vision, and spoke English as their first language (or had acquired English as a second language by age five). Current drug use excepting nicotine was cause for exclusion, as were history of alcohol-related liver disease, epilepsy, head trauma resulting in loss of consciousness for 15 minutes or more, HIV, schizophrenia, or metal implants.

### 2.2 Neuropsychological Assessment

Neuropsychological testing was conducted at the Department of Veterans Affairs (VA) Boston Healthcare System facility prior to scanning. Participants completed a medical history interview, vision test, handedness questionnaire (Briggs & Nebes, 1975), and a battery of tests as described below. All subjects were screened using the Hamilton Rating Scale for Depression (Hamilton, 1960) and the Diagnostic Interview Schedule for the DSM-IV (Robins *et al*., 2000). The majority of participants also were administered the Wechsler Adult Intelligence Scale (WAIS-III) and the Wechsler Memory Scale (WMS-III) (Wechsler, 1997). Four participants (two ALCw and two ALCm) received the WAIS-IV and WMS-IV (Holdnack & Drozdick, 2010), and WMS-III scores were not obtained from one ALCm. The scores for these participants were adjusted to account for differences in scoring outcomes relative to the earlier versions of the scales. Because craving for the rewarding effects of alcohol is known to serve as a trigger for relapse in those recovering from AUD (Schneider *et al*., 2001), and alcohol cue exposure in particular is known to be related to relapse (Lubman, 2007), all participants were administered the Penn Alcohol Craving Scale (Flannery *et al*., 1999) immediately before and approximately two weeks following the scan to assess any changes in alcohol craving patterns.

### 2.3 Alcohol Screening

The ALC participants met criteria for alcohol abuse or dependence, and consumed 21 or more alcoholic drinks per week for five or more years. Extent of alcohol use was assessed by calculating Quantity Frequency Index (QFI) scores (Cahalan *et al*., 1969). QFI scores approximate the number of drinks consumed per day, and take into consideration the amount, type, and frequency of alcohol consumption either over the last six months (NC participants), or over the six months preceding cessation of drinking (ALC participants), and yields an estimate of ounces of ethanol per day. To remove the influence of current alcohol abuse, ALC participants must have been abstinent for at least four weeks before the scan date to be included. The ALC participants did not display symptoms of Korsakoff’s Syndrome nor dementia (Oscar-Berman & Maleki, 2019). Potential NC participants who had consumed 15-20 drinks per week for any length of time or who engaged in binge drinking were disqualified.

### 2.4 Functional Imaging Task

All participants were given a delayed matching-to-sample memory task (Dolcos & McCarthy, 2006) in a magnetic resonance imaging (MRI) scanner, whereby they were asked to encode two faces that both had one of three emotional valences: positive, neutral, or negative (Figure 1). The face stimuli were shown in grayscale and were taken from a set of faces used in a previous study (Marinkovic *et al*., 2009). These faces were displayed simultaneously for three seconds, followed by an asterisk (*) for one second. Subjects were asked to maintain these faces in memory while a colored distractor stimulus was shown. On different trials, the distractor stimulus was either a picture of an alcoholic beverage (*alcbev*; beer, wine, liquor, or mixed drink), a picture of a nonalcoholic beverage (*nonalcbev*; water, juice, milk, soda, coffee, tea, etc.), or a scrambled nonsense picture (*scrambled*). Alcoholic and nonalcoholic beverage pictures were a combination of images used with permission from the Normative Appetitive Picture System (NAPS) (Stritzke *et al*., 2004), and other previously published works on alcohol cues (Grüsser *et al*., 2000; Wrase *et al*., 2002; Myrick *et al*., 2004). Additional distractor images were modified from digital photographs taken at bars, liquor stores, and convenience stores. The scrambled images were created by inverting half the alcoholic and half the nonalcoholic beverage images and distorting them until they were not recognizable as any particular object, while preserving a match of primary visual characteristics. Each distractor picture was shown for three seconds, followed by an asterisk (*) for one second. Following the distractor picture, a single probe face was shown for two seconds, and the participants were instructed to report whether this face was one of the two faces they had just seen.

**Figure 1.**
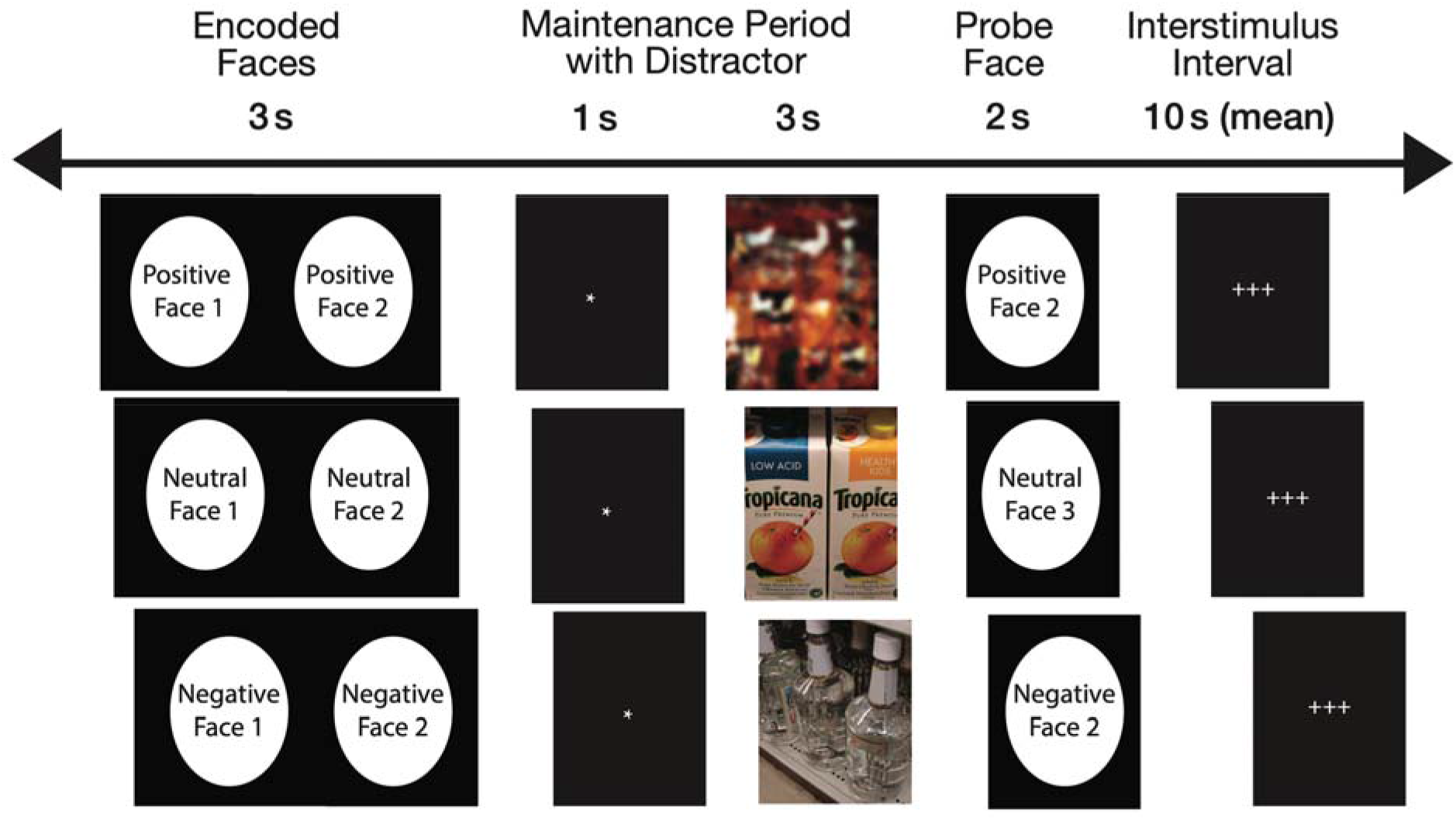
Task presented during functional neuroimaging. Two faces were presented simultaneously for three seconds, followed by an asterisk for one second. Next, a distractor was presented for three seconds. The probe face immediately followed, during which the subjects had been trained to respond with a button press with either their index or middle finger to indicate whether the probe face matched the encoded face. Three crosses served as the inter-trial interval, which lasted from 2 to 30 seconds (mean 10 seconds). A total of 162 trials were presented. While the faces in this figure have been blurred to mask the identities of the individuals, the research participants saw the original unblurred photographs.

Each trial was 10 seconds in length, and was followed by a variable delay period (with a mean duration of 10 seconds, ranging from 2-22 seconds) during which the subject saw a set of crosshairs (+++) serving as a visual fixation. The task was divided into nine runs, each of which contained 18 trials. There were nine trial types made up of each combination of face valence and distractor type (*e.g.*, positive faces followed by alcohol distractor). Each emotion-distractor combination appeared twice per run. The stimulus order and variable inter-trial intervals were determined using *optseq2* (http://surfer.nmr.mgh.harvard.edu/optseq), which optimizes statistical efficiency and hemodynamic response estimate accuracy for event-related experimental designs (Dale, 1999). In total, there were 54 trials for each distractor type (combined across facial expressions) and each face valence (combined across distractor types), for a total of 162 trials across the entire scan. The stimulus faces were balanced to contain 50% male and 50% female faces. Within a trial, the two encoded faces and probe face were matched on emotional expression and gender. This way, on match trials the probe facial image was identical to one of the encoded images, and on mismatch trials the facial identity changed but the emotional expression and gender did not.

The probe face matched one of the encoded faces on 50% of the trials, and match/mismatch trials appeared in a randomized order within each run. Responses were made by pressing one of two buttons with the index finger (match) or middle finger (mismatch) of the right hand. Participants were instructed to respond as quickly as possible without sacrificing accuracy. Additionally, participants could immediately correct a response by pressing the opposite button. To ensure the distractor images were viewed by all participants, they were told that it was necessary to pay attention to the pictures shown in between the faces on each trial, as they would be questioned about those images following the scan.

### 2.5 Behavioral Task Analyses

Responses to the face memory task were analyzed on the first level (individual subjects) using custom Excel templates. Trials were organized by Distractor type, facial Emotion, and Face Gender. Each trial was scored as correct, incorrect, or miss (*i.e.*, no response). When more than one response was made to a trial, the last response type (*i.e.*, yes/match or no/nonmatch) was accepted as the final answer, provided that the final response was at least 200 ms after the preceding response and no more than 10 s following the preceding response. When a single response was recorded and the reaction time was less than 200 ms, the trial was scored as a miss. For each participant, a mean overall reaction time (regardless of trial type) was calculated. Reaction times that exceeded three standard deviations from this mean were excluded from reaction time calculations by trial type. Participants’ patterns of responses were analyzed for consecutive misses to assure that they remained awake throughout the task. Three separate runs were identified, each in a different participant, wherein greater than five consecutive trials were missed; these runs were excluded from behavioral analyses.

Second level (group) effects on percent correct and reaction time (correct trials) were investigated using SPSS Version 17.0 (IBM, Chicago, IL, USA). Repeated-measures analyses of variance (ANOVA) were carried out with between-subjects factors of Group (ALC or NC) and Gender (female participant or male participant) and within-subjects factors of Distractor (alcbev, nonalcbev, or scrambled), Emotion (positive, negative, or neutral), and Face Gender (female face or male face).

### 2.6 Image Acquisition

Imaging was conducted at the Massachusetts General Hospital’s Athinoula A. Martinos Center for Biomedical Imaging in Charlestown, MA. Data were acquired on a 3 Tesla Siemens (Erlangen, Germany) MAGNETOM Trio Tim MRI scanner with a 12-channel head coil. Sagittal T1-weighted MP-RAGE scans (TR = 2530 ms, TE = 3.39 ms, flip angle = 7°, FOV = 256 mm, slice thickness = 1.33 mm, slices = 128, matrix = 256 x 192) were collected for all subjects. For most participants, two such volumes were collected and averaged to aid in motion correction. An auto-align localizer was employed to adjust the acquired slices such that they ran parallel to an imaginary plane between the anterior and posterior commissures. Echo planar functional MRI blood oxygen level dependent (BOLD) scans were collected axially with 5 mm slice thickness and 3.125 x 3.125 mm in-plane resolution (64 x 64 matrix), allowing for whole brain coverage (32 interleaved slices, TR = 2 s, TE = 30 ms, flip angle = 90°). The event-related design included 18 trials per run with a total of nine runs. Within each six-minute run, 180 T_2_*-weighted volumes were collected. Functional volumes were auto-aligned to the anterior/posterior commissure line to ensure a similar slice prescription was employed across participants. Prospective Acquisition Correction (3D-PACE) was applied during collection of the functional volumes to minimize the influence of participants’ body motion (Thesen *et al*., 2000). An IBM ThinkPad (Windows XP) running Presentation version 11.2 (NeuroBehavioral Systems, Albany, CA) software was used for visual presentation of the experimental stimuli and collection of participants’ responses. Stimuli were back-projected onto a screen at the back of the scanner bore and were viewed by the participants through a mirror mounted to the head coil. All participants wore earplugs to attenuate scanner noise.

### 2.7 Structural Image Processing

Structural MPRAGE image analyses were performed for all participant data using the FreeSurfer (version 4.5.0) image analysis suite (http://surfer.nmr.mgh.harvard.edu). A multi-stage cortical surface reconstruction process was run on the two collected T1-weighted MP-RAGE scans (Dale *et al*., 1999), starting with motion correction, intensity normalization (Sled *et al*., 1998), Talairach registration (Talairach & Tournoux, 1988), skull stripping (Ségonne *et al*., 2004), and segmentation (Fischl *et al*., 2002) of white matter, gray matter, and ventricles. Subsequently, boundaries were calculated delineating where gray and white matter meet, and where gray matter adjoins cerebrospinal fluid (“pial surfaces”) based on maximal shifts in image intensity between tissue types. These boundaries, as well as the subcortical segmentations, were visually inspected on each coronal slice for every subject, and manual interventions (*e.g.*, white matter volume corrections) were made when needed. The surface boundaries were used to generate computationally inflated two-dimensional cortical surface models, which allowed individual subjects to be registered to a spherical atlas by utilizing each subject’s cortical folding patterns. This registration was used to align the cortical geometry of all subjects within a group. Creation of these cortical surface models allowed improved data visualization as well as improved accuracy of within-group co-registration relative to an affine morph procedure (Fischl *et al*., 1999). The cortical surface models were employed in an automated parcellation procedure that divides the surface into subregions based on gyral and sulcal anatomy. The Destrieux atlas parcellation for FreeSurfer (Destrieux *et al*., 2010) was used to define anatomical ROI in the functional analyses.

### 2.8 Functional Image Processing and Statistical Analyses

Effects of Group, Gender, Distractor, and Emotion on the BOLD signal were evaluated using both a whole-brain cluster analysis as well as ROI analyses. Processing of the functional data was performed using the FreeSurfer Functional Analysis Stream (FS-FAST) version 5.3, SPSS Version 17.0, and Matlab 7.4.0.

#### 2.8.1 First-Level Functional Analyses

Preprocessing of the functional images for first-level (individual subject) FS-FAST analysis included motion correction, intensity normalization (Sled *et al*., 1998), and spatial smoothing with a 5-mm Gaussian convolution kernel at full-width half-maximum. Trials were first combined across runs by distractor-emotional face valence pairs (i.e., alcbev-positive, alcbev-negative, alcbev-neutral, nonalcbev-positive, nonalcbev-negative, nonalcbev-neutral, scrambled-positive, scrambled-negative, scrambled-neutral) and then collapsed across emotional valence. The BOLD response was estimated using a Finite Impulse Response (FIR) model, which allows for estimation of the time course of activity (percent signal change for a given condition) within a vertex or ROI for the entire trial period. For each condition, estimates of signal intensity were calculated for 2 pre-trial and 10 post-trial onset TRs, for a total analysis window of 24 seconds. Motion correction parameters calculated during alignment of the functional images were entered into the analysis as external regressors. Alignment of the T2*-weighted functional images with T1-weighted structural volumes was accomplished through an automated boundary-based registration procedure (Greve & Fischl, 2009). These automated alignments were manually inspected to ensure accuracy.

Statistical maps were generated for each of the 84 individual subjects for contrasts between experimental conditions. Three contrasts were used to identify DMN regions (as described below); they were made between distractor types and fixation: (1) alcbev vs. fixation, (2) nonalcbev vs. fixation, and (3) scrambled vs. fixation. Another three contrasts were used to assess cue responsivity: (1) alcbev vs. nonalcbev, (2) alcbev vs. scrambled, (3) nonalcbev vs. scrambled. Analyses of each of these contrasts included removal of prestimulus differences between the contrasted conditions by averaging the first three time points (two pre-trial onset and one post-trial onset) for each condition and subtracting this mean from each time point for that condition. Time points summed for inclusion in each contrast were chosen to reflect peak stimulus-related activity: FIR estimates of hemodynamic responses to the distractors were analyzed using a mean of the five TRs collected during the time period of 2-12 seconds post distractor onset. Since the distractor is shown 4 seconds after the trial onset, the analysis window is 6-16 seconds following trial onset (time points 3 through 8).

#### 2.8.2 Cortical Surface Cluster Analyses

We investigated cue-related brain activation in two separate cortical brain networks: (1) cue reactivity in DMN regions, and (2) cue reactivity in task-regions. The brain network that was more active during presentation of the fixation cue than during the distractor images was used as our measure of DMN regions (fixation-regions). The network that was more active during the presentation of distractor images than during presentation of the fixation stimulus was used as our measure of the task-regions. In what follows, we first describe masking procedures and analyses we used to separate the networks.

The *t*-statistic maps for each condition vs. fixation were thresholded at *p* < 0.05 vertex-wise and were used to generate binary masks (Figure 2) separating fixation-regions and task-regions, thereby forming the six masks: alcbev greater than (or less than) fixation, nonalcbev greater than (or less than) fixation, scrambled greater than (or less than) fixation. These masks were used to separate the between-distractor analyses (described below).

**Figure 2.**
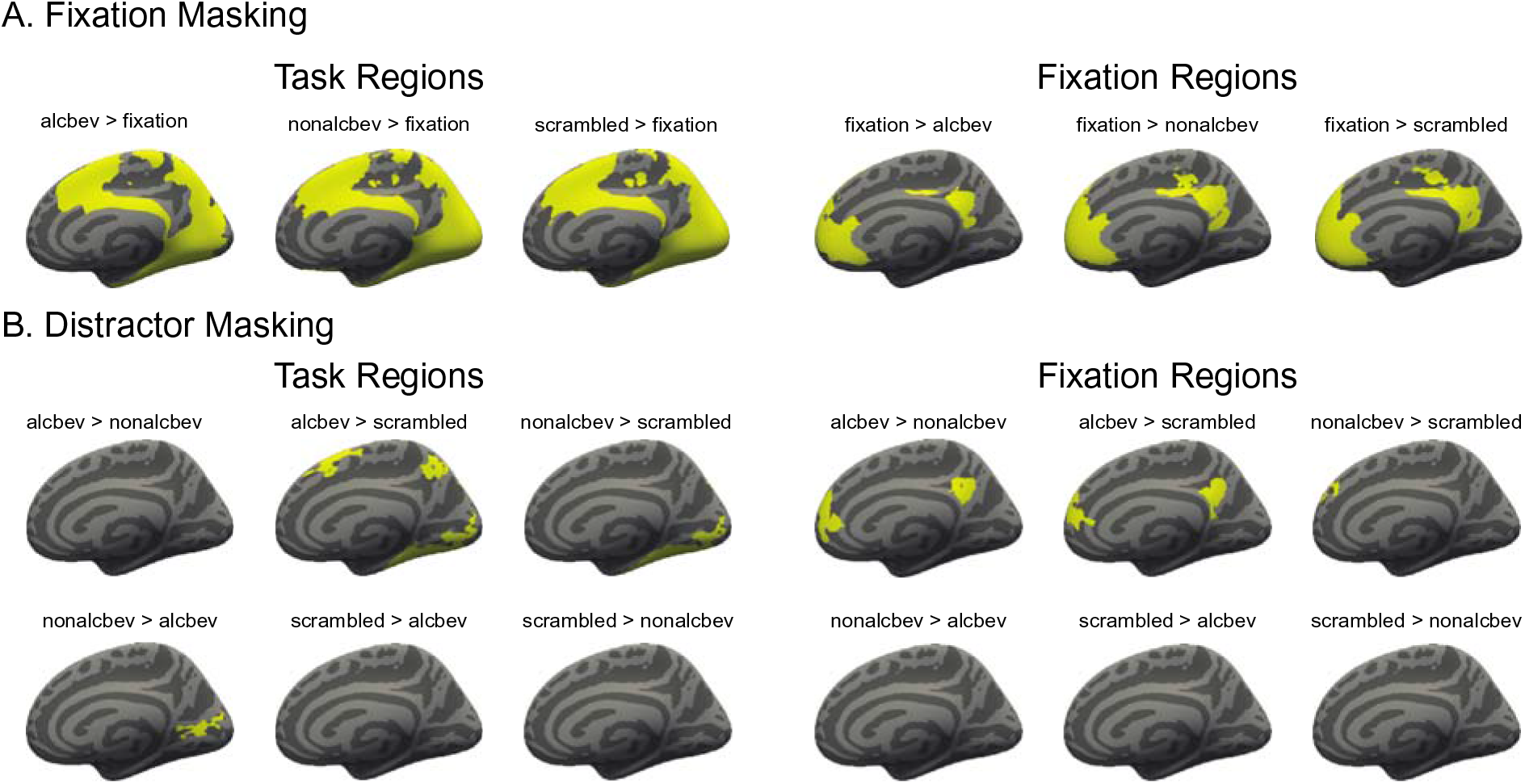
Analyses conducted with masked fMRI data for cortex. The top set of brains represents analyses conducted for cortical regions in which network masking (for task and fixation-regions) were performed. The bottom set of brains represents analyses conducted for cortical regions in which distractor masking (for task and fixation-regions) were performed.

Second-level (group) analyses on cortical regions were accomplished using FS-FAST, a surface-based morphing procedure for intersubject alignment and statistics (Fischl *et al*., 1999). Group-averaged signal intensities during each experimental condition (alcbev, nonalcbev, scrambled) relative to fixation were calculated using the general linear model in spherical space for cortical regions, and were mapped onto the canonical cortical surface *fsaverage*, generating group-level weighted random-effects *t*-statistic maps masked to include only the cortex. Weighted random effects models were employed to reduce noise by taking into account individual subject variance. A 5 mm smoothing kernel (full-width half-maximum) was employed for all group and intergroup maps. Cluster correction on maps showing activity for each distractor condition vs. fixation was applied using FS-FAST Monte Carlo simulation with a clusterwise threshold of *p* < 0.05 corrected for three spaces (left hemisphere cortical, right hemisphere cortical, and subcortical). Cortical surface cluster regions were identified by the location of each cluster’s peak vertex on the cortical surface according to the Desikan-Killiany atlas (Desikan *et al*., 2006).

When we examined brain activation differences between the distractor types, we used three contrasts: alcbev vs. nonalcbev, alcbev vs. scrambled, and nonalcbev vs. scrambled. We investigated each direction of these contrasts separately. For example, brain regions with higher activation for alcbev than nonalcbev would be analyzed separately from those regions with higher activation for nonalcbev than alcbev.

Intergroup comparison *t*-statistic maps were generated using FS-FAST by comparing activation levels of all of the ALC participants with levels of all of the NC participants. Additionally, Group-by-Gender interaction maps for each contrast were calculated.

#### 2.8.3 Region of Interest Analyses

The anatomically-defined ROI for the distractor analyses included areas hypothesized *a priori* to be implicated in alcohol craving, distractor interference, and working memory for emotional faces, as described in the Introduction. These were DLPFC, VLPFC, OFC, insular cortex, parahippocampal gyrus, hippocampus, amygdala, fusiform, ACC, and the multi-regional EROS (Makris *et al*., 2008; Sawyer *et al*., 2017). Left and right hemisphere regions were analyzed as separate ROI.

Statistical preprocessing and time course visualization of ROI data were performed using scripts written for Matlab version 7.4.0. Signal intensity for each region was averaged across all vertices (for surface-based ROI) or voxels (for volume-based ROI) included in the region for each condition on the individual participant level. To compute percent signal change for each participant within an ROI, signal estimate per condition and time point was divided by the average baseline activity for that participant. Time courses were normalized at the individual subject level for each condition by taking the mean of the first three time points (two pre-trial and one post-trial onset) and subtracting this mean from each time point. Group and Group-by-Gender averages of the normalized time courses were computed for each condition, and were visualized by plotting the percent signal change for each condition at each time point (*i.e.*, TR) of the trial.

For the distractor ROI analyses, percent signal changes of the BOLD signal within each ROI for the time window from 2 to 12 sec after distractor onset were entered as dependent variables into repeated-measures ANOVA models with between-group factors of Group (ALC or NC) and Gender (men or women) and within-subjects factor of Distractor type (alcbev, nonalcbev, or scrambled).

## 3. Results

### 3.1 Research Participant Characteristics

Table 1 summarizes means, standard deviations, and ranges of participant demographics, drinking variables, and IQ and memory test scores.

The ALC and NC groups did not differ significantly by age. Although the NCw were older than the NCm, controls did not differ significantly from their respective ALC counterparts by age. ALCm had on average one year less education relative to NCm. Groups did not differ significantly on WAIS-III Full Scale IQ scores. While ALCw had higher Hamilton Rating Scale for Depression scores than AUDm, the average scores for all four subgroups (ALCm, ALCw, NCm, and NCw) were low (all means below 5, whereas mild depression threshold is 8), so depression likely contributed little to our observed gender differences.

By definition, the ALC group had longer durations of heavy drinking than the NC group. The ALCm on average drank heavily for six years more than did the ALCw, and showed a trend toward drinking larger average daily quantities (QFI, *F_1,40_* = 3.66, *p* = 0.06). Five NCm and two NCw reported being lifetime abstainers (and as such did not have relevant length of sobriety values). The ALC group reported higher levels of craving for alcohol than the NC group on the PACS administered immediately prior to the scan; ALCm and ALCw did not differ on reported level of pre-scan alcohol craving. Eighty-one of the 84 participants were reached approximately two weeks after their scan date to be reassessed on alcohol craving level. One ALCw, one ALCm, and one NCw could not be reached for follow up assessment on PACS scores. Neither the ALC group nor the NC group displayed an increase in alcohol craving (*i.e.*, a significant change in PACS scores) from the assessment on their scan date to their follow up assessment.

### 3.2 Behavioral Results

Measures of participant performance on the face memory task were calculated for overall performance and for performance by each Distractor type and facial Emotion. Means, standard deviations, and ranges are reported for percent correct responses and reaction times in Table A1 and Table A2, respectively.

A significant Group-by-Gender interaction was found for accuracy (*F_1,80_* = 6.880, *p* = 0.01, Figure A1 and Table A1). The significant interaction indicated that the better performance for the ALCw than the ALCm was larger than the difference between NCw and NCm. Accuracy and reaction times did not vary significantly as a function of the Distractor, nor were there any significant interactions of Distractor with Group or Gender (all *p* > 0.05). The main effect of Emotion was significant for percent correct responses, wherein ALC and NC participants alike performed better on both positive and negative faces relative to neutral faces. Performance on positive and negative faces did not differ significantly. The effect of Emotion on percent correct responses did not vary as a function of Group or Gender (all *p* > 0.05). Regarding reaction time and Emotion, participants responded more quickly to positive face trials relative to neutral face trials; this effect did not vary by Group nor by Gender. The main effect of Face Gender on percent correct responses also was significant (*F_1,80_* = 6.80, *p* = 0.01), with the overall performance being better for male than for female faces. The effects of Group and Gender on percent correct responses and reaction times for Face Gender were not significant (all *p* > 0.05).

### 3.3 fMRI BOLD Effects

Effects of the distractors on the BOLD signal were assessed using group and intergroup cluster analyses for cerebral cortex, along with a-priori analyses of anatomical ROI that had been implicated by the literature. Below, we report group analyses of fixation contrasts and between-distractor conditions, followed by intergroup analyses of the same contrasts.

#### 3.3.1 Cortical Cluster Analyses of Distractor Effects

Analyses of task contrasts revealed broadly similar activation patterns for the ALCw, ALCm, NCw, and NCm groups. During fixation, regions involved in the DMN (the anatomical network described in the Introduction) were significantly more active than during the presentation of distractor images. We refer to those more active regions as fixation-regions. As detailed in the Methods, these regions were masked and examined separately for subsequent analyses of contrasts between distractor types. Identical analyses were then performed for the task-regions. Significant clusters for between-distractor contrasts can be seen for fixation-regions first (Figure 3 and Figure A3), and then for task-regions (Figure 4 and Figure A4).

**Figure 3.**
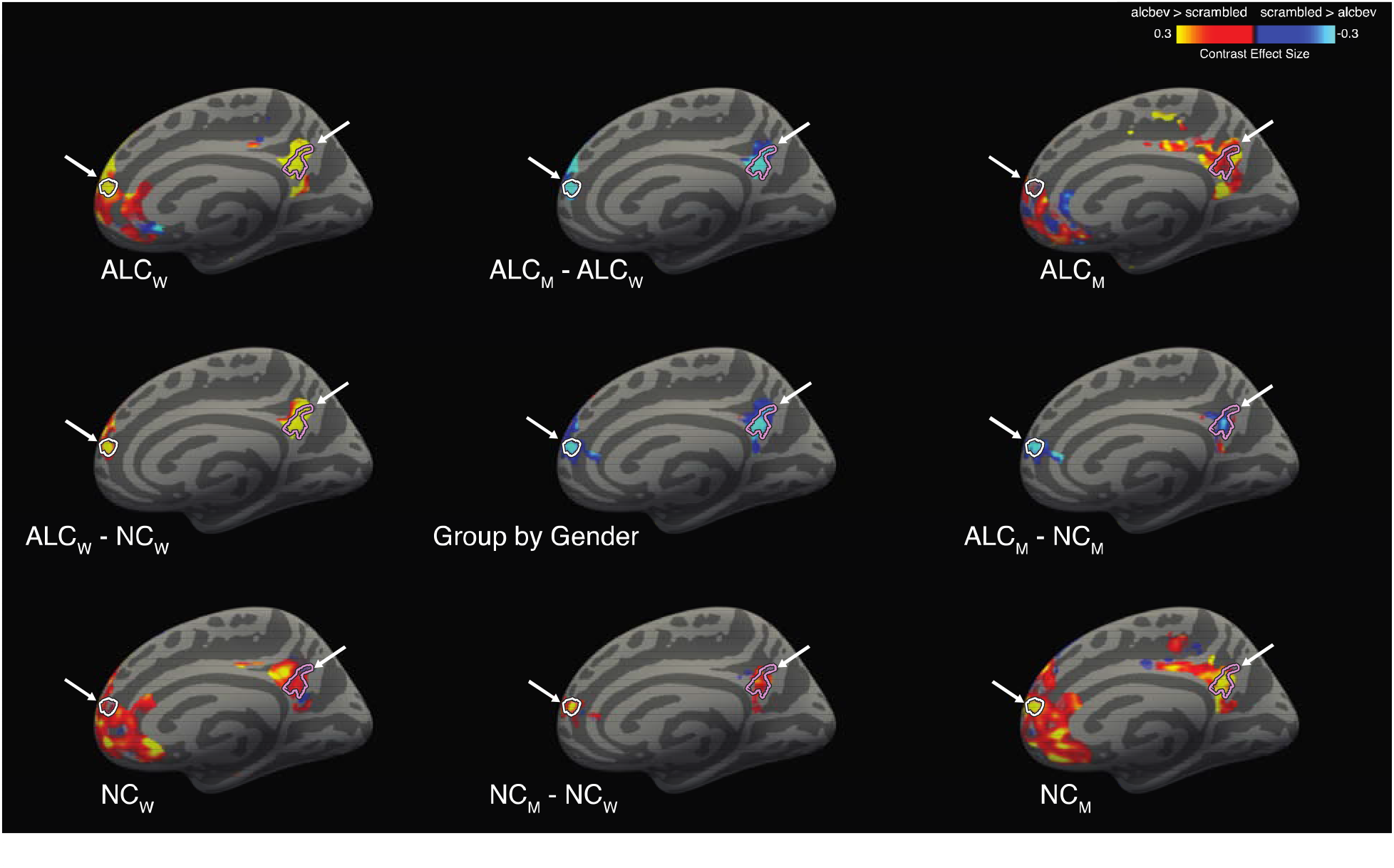
Cortical clusters significant for Group-by-Gender interactions for the alcbev vs. scrambled contrast within fixation-regions (right medial view) A significant Group-by-Gender interaction revealed several clusters (see Table A4), two of which are indicated by arrows on the medial surface of the right hemisphere, with cluster outlines overlaid on contrast values between alcbev and scrambled distractors. Group mean contrast values (for alcbev vs. scrambled within fixation-regions) are displayed in the four brain images located in the corners of the figure, and group comparisons are indicated by minus signs. Abbreviations: ALCm = Alcoholic men; ALCw = Alcoholic women; NCm = Nonalcoholic men; NCw = Nonalcoholic women.

**Figure 4.**
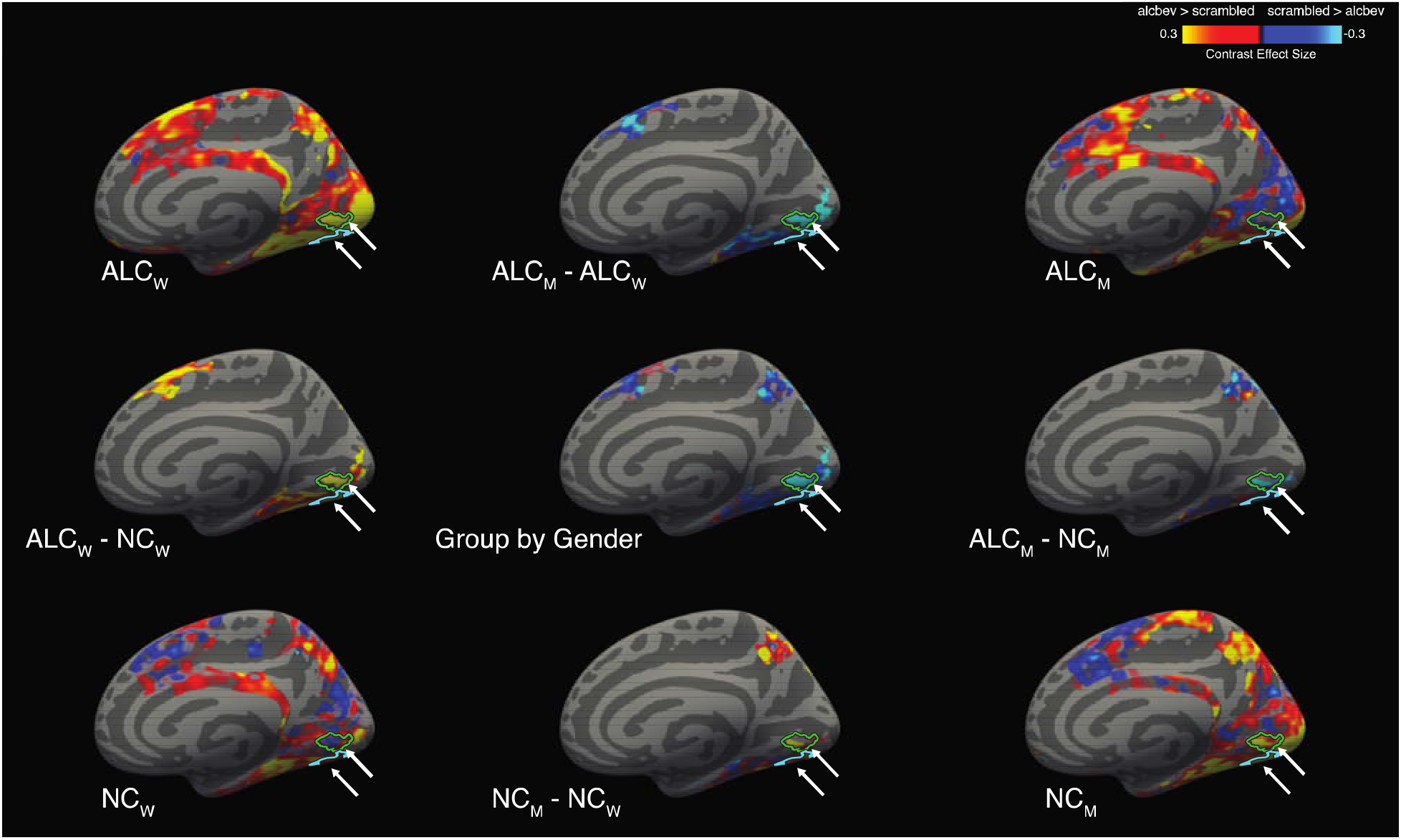
Cortical clusters significant for Group-by-Gender interactions, for the alcbev vs. scrambled contrast within task-regions (right medial view) A significant Group-by-Gender interaction revealed several clusters (see Table A4), two of which are indicated by arrows on the medial surface of the right hemisphere, with cluster outlines overlaid on contrast values between alcbev and scrambled distractors. Group mean contrast values (for alcbev vs. scrambled within task-regions) are displayed in the four brain images located in the corners of the figure, and group comparisons are indicated by minus signs. Abbreviations: ALCm = Alcoholic men; ALCw = Alcoholic women; NCm = Nonalcoholic men; NCw = Nonalcoholic women.

All four groups had more brain activity (Table A3) in response to alcbev than scrambled distractors in the four main fixation-regions (the anterior and posterior medial hub regions, the temporal parietal junction, and the middle temporal gyrus), while the nonalcbev vs. scrambled contrast was less consistent. The alcbev vs. nonalcbev contrast generally indicated higher activation for the alcbev than nonalcbev. For task-regions, alcbev and nonalcbev elicited higher activation than scrambled in the occipital lobe and adjoining visual areas in temporal and parietal cortex (Table A3, Figure A4).

#### 3.3.2 Distractor Intergroup Cluster Analyses

The pattern of results indicated that ALCw and NCm had strong activation contrasts (beverages > scrambled) in visual areas and the medial DMN regions, especially the posterior hub. The ALCw had greater activation contrast than NCw, while ALCm had lower activation contrast than NCm. Table 2 summarizes the regions where significant Group-by-Gender interactions were found for the distractor contrasts, i.e., for alcbev vs. nonalcbev, alcbev vs. scrambled, and nonalcbev vs. scrambled. Table A4 provides all significant Group-by-Gender interactions, and Figure 3, Figure 4, Figure A3, Figure A4 illustrate alcbev vs. scrambled contrasts. In total, we observed 22 clusters where the Group-by-Gender interaction was statistically significant: Seven in fixation-regions and 15 in task-regions.

**Table 2.**
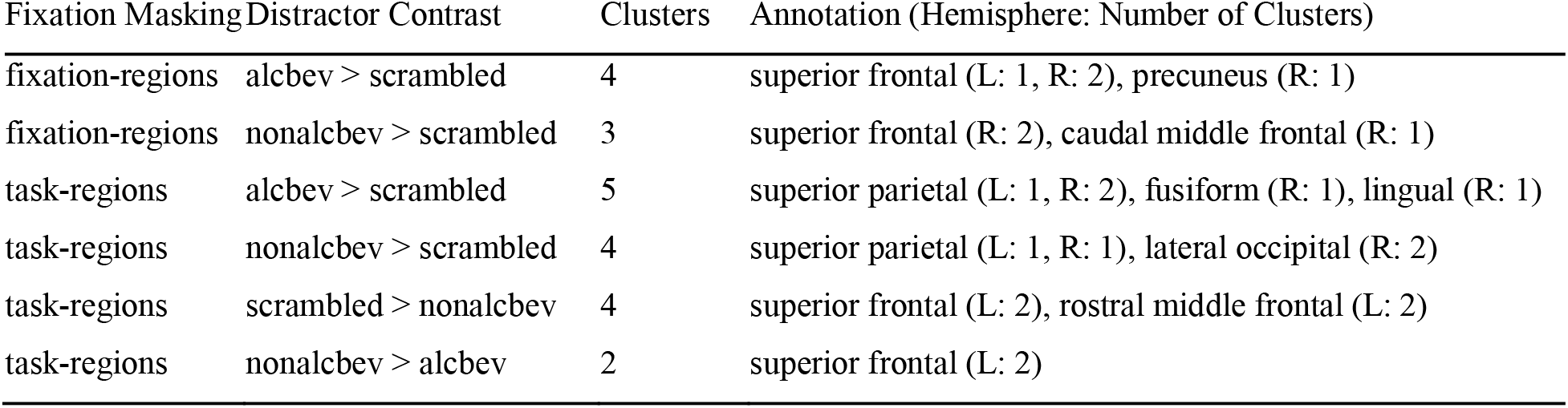
Group-by-gender cortical cluster summary. Annotations (using the Desikan-Killiany atlas) are shown for each of the 22 clusters with significant Group-by-Gender interactions in distractor contrasts. The Fixation Masking column refers to the separate analyses conducted for fixation-regions and task-regions as shown in Figure 2. The clusters reported can be understood to span multiple functional regions [71]. That is, they are not limited to a single region, as reported by the maximal vertex or voxel. Abbreviations: alcbev = alcoholic beverages; nonalcbev = nonalcoholic beverages; L = left hemisphere; R = right hemisphere. See Table A4 for detailed cluster information. Note: for the fixation-regions, superior frontal and caudal middle frontal are part of the anterior hub; and precuneus is part of the posterior hub.

For the seven clusters in fixation-regions, six were in the anterior hub and one was in the posterior hub. For alcbev > scrambled, we observed one cluster in the posterior hub and three in the anterior hub. For three of these four clusters, ALCw and NCm had the strongest contrasts; in the fourth cluster, ALCw had the strongest contrast, whereas ALCm had the weakest. The remaining three clusters were found for nonalcbev > scrambled. As with the alcbev contrasts, ALCw and NCm had the strongest contrasts. All three clusters were in the right hemisphere of the anterior hub. Thus, the overall pattern consistent among the seven clusters was as follows: NCm had stronger contrast (beverage > scrambled) than ALCm, while muted or opposite direction comparison was observed for the women.

Of the 15 task-regions clusters, two were found for nonalcbev > alcbev (left superior frontal cortex). In both clusters, NCw had higher activation to nonalcoholic beverages, while NCm had higher activation to alcoholic beverages. In the other 13 clusters, the strongest contrasts were found for NCm and ALCw. Of those, the significant interactions were found in visual regions where nonalcbev > scrambled, while significant interactions in frontal regions were found for scrambled > nonalcbev.

Setting aside gender, analyses of activation levels comparing ALC and NC groups revealed 15 cortical clusters with significantly greater contrast levels for the ALC group than for the NC group (Table A5). Two of the clusters were in the DMN: one alcbev-region (temporoparietal junction), and one nonalcbev-region (posterior hub). The other 13, all alcbev-regions, were in task-regions located throughout the cortex: 7 frontal, 3 temporal, 2 parietal, and 1 occipital.

#### 3.3.3 Distractor Region of Interest Analyses

Results of ANOVAs examining between-subjects effects of Group and Gender and within-subjects effects of Distractor type on BOLD percent signal change within each ROI are summarized in Table 3. Reported means and standard deviations represent the percent signal change across each ROI (unmasked) for the time period of 2 to 12 seconds post-distractor stimulus onset. These anatomically-defined ROI included regions in EROS areas (Makris *et al*., 2008; Sawyer *et al*., 2017), as well as in regions associated with face memory maintenance and distractor interference, as described in the Introduction and Methods.

**Table 3.**
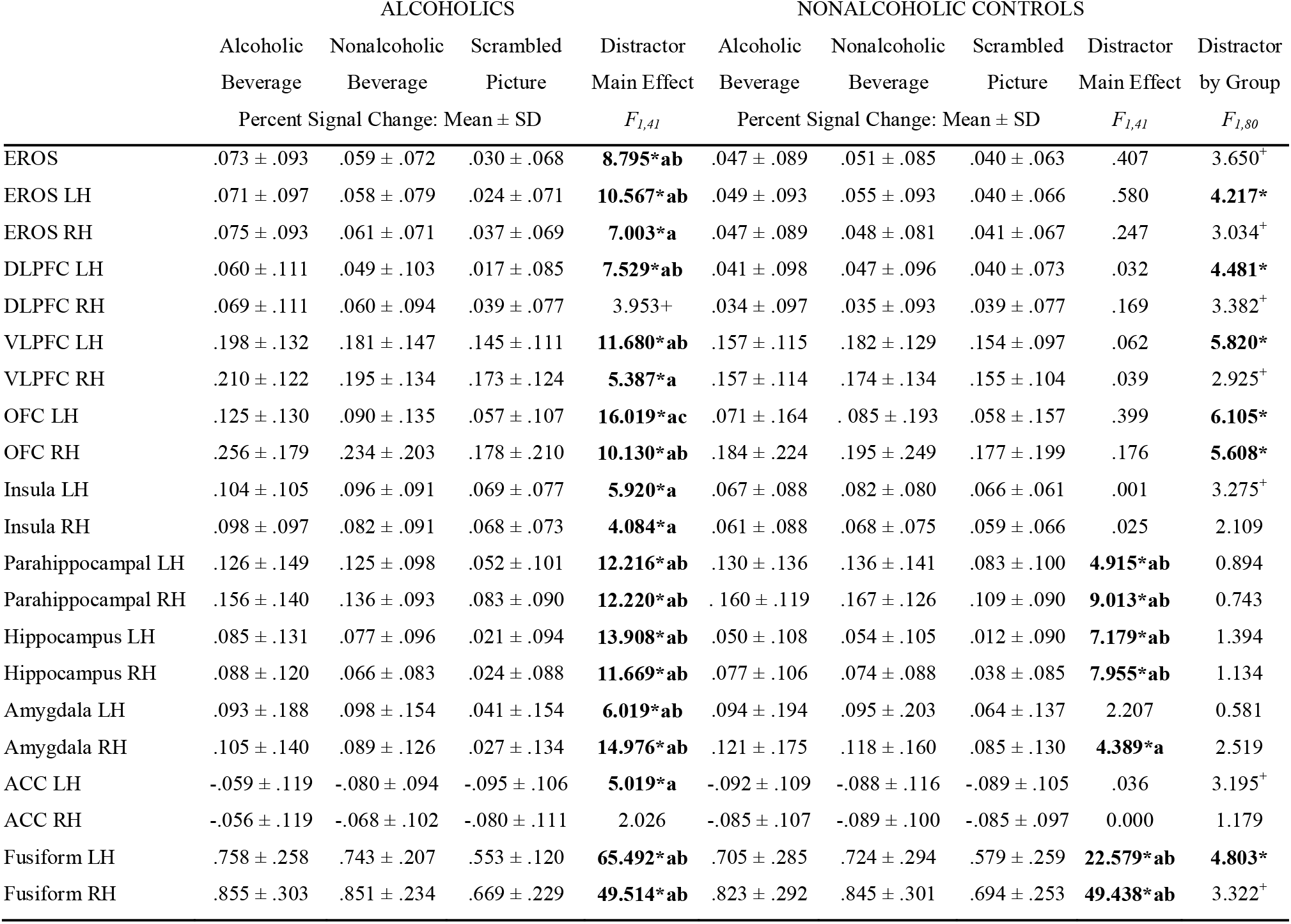
Percent signal change for each distractor type by group, for a-priori regions. Distractor by Group F-values are reported from a full factorial ANOVA model with between-groups factors of Group and Gender and within-groups factor of Distractor type. Distractor main effect F-values within each group are reported from ANOVA model including within-subjects factor of Distractor type. Abbreviations: EROS = Extended Reward and Oversight System; DLPFC = Dorsolateral Prefrontal Cortex; VLPFC = Ventrolateral Prefrontal Cortex; OFC = Orbitofrontal Cortex; ACC = Anterior Cingulate Cortex; LH = Left Hemisphere; RH = Right Hemisphere a. Alcoholic Beverage > Scrambled Picture, *p* < .05 b. Nonalcoholic Beverage > Scrambled Picture, *p* < .05 c. Alcoholic Beverage > Nonalcoholic Beverage, *p* < .05 \**p* < .05 +.05 <*p* <.10

|  | ALCOHOLICS |  |  |  | NONALCOHOLIC CONTROLS |  |  |  |  |
| --- | --- | --- | --- | --- | --- | --- | --- | --- | --- |
|  | Alcoholic | Nonalcoholic | Scrambled | Distractor | Alcoholic | Nonalcoholic | Scrambled | Distractor | Distractor |
|  | Beverage | Beverage | Picture | Main Effect | Beverage | Beverage | Picture | Main Effect | by Group |
| | Percent Signal Change: Mean ± SD | | | $F_{1,41}$ | Percent Signal Change: Mean ± SD | | | $F_{1,41}$ | $F_{1,80}$ |
| EROS | .073 ± .093 | .059 ± .072 | .030 ± .068 | **8.795\*ab** | .047 ± .089 | .051 ± .085 | .040 ± .063 | .407 | 3.650<sup>+</sup> |
| EROS LH | .071 ± .097 | .058 ± .079 | .024 ± .071 | **10.567\*ab** | .049 ± .093 | .055 ± .093 | .040 ± .066 | .580 | **4.217\*** |
| EROS RH | .075 ± .093 | .061 ± .071 | .037 ± .069 | **7.003\*a** | .047 ± .089 | .048 ± .081 | .041 ± .067 | .247 | 3.034<sup>+</sup> |
| DLPFC LH | .060 ± .111 | .049 ± .103 | .017 ± .085 | **7.529\*ab** | .041 ± .098 | .047 ± .096 | .040 ± .073 | .032 | **4.481\*** |
| DLPFC RH | .069 ± .111 | .060 ± .094 | .039 ± .077 | 3.953<sup>+</sup> | .034 ± .097 | .035 ± .093 | .039 ± .077 | .169 | 3.382<sup>+</sup> |
| VLPFC LH | .198 ± .132 | .181 ± .147 | .145 ± .111 | **11.680\*ab** | .157 ± .115 | .182 ± .129 | .154 ± .097 | .062 | **5.820\*** |
| VLPFC RH | .210 ± .122 | .195 ± .134 | .173 ± .124 | **5.387\*a** | .157 ± .114 | .174 ± .134 | .155 ± .104 | .039 | 2.925<sup>+</sup> |
| OFC LH | .125 ± .130 | .090 ± .135 | .057 ± .107 | **16.019\*ac** | .071 ± .164 | .085 ± .193 | .058 ± .157 | .399 | **6.105\*** |
| OFC RH | .256 ± .179 | .234 ± .203 | .178 ± .210 | **10.130\*ab** | .184 ± .224 | .195 ± .249 | .177 ± .199 | .176 | **5.608\*** |
| Insula LH | .104 ± .105 | .096 ± .091 | .069 ± .077 | **5.920\*a** | .067 ± .088 | .082 ± .080 | .066 ± .061 | .001 | 3.275<sup>+</sup> |
| Insula RH | .098 ± .097 | .082 ± .091 | .068 ± .073 | **4.084\*a** | .061 ± .088 | .068 ± .075 | .059 ± .066 | .025 | 2.109 |
| Parahippocampal LH | .126 ± .149 | .125 ± .098 | .052 ± .101 | **12.216\*ab** | .130 ± .136 | .136 ± .141 | .083 ± .100 | **4.915\*ab** | 0.894 |
| Parahippocampal RH | .156 ± .140 | .136 ± .093 | .083 ± .090 | **12.220\*ab** | .160 ± .119 | .167 ± .126 | .109 ± .090 | **9.013\*ab** | 0.743 |
| Hippocampus LH | .085 ± .131 | .077 ± .096 | .021 ± .094 | **13.908\*ab** | .050 ± .108 | .054 ± .105 | .012 ± .090 | **7.179\*ab** | 1.394 |
| Hippocampus RH | .088 ± .120 | .066 ± .083 | .024 ± .088 | **11.669\*ab** | .077 ± .106 | .074 ± .088 | .038 ± .085 | **7.955\*ab** | 1.134 |
| Amygdala LH | .093 ± .188 | .098 ± .154 | .041 ± .154 | **6.019\*ab** | .094 ± .194 | .095 ± .203 | .064 ± .137 | 2.207 | 0.581 |
| Amygdala RH | .105 ± .140 | .089 ± .126 | .027 ± .134 | **14.976\*ab** | .121 ± .175 | .118 ± .160 | .085 ± .130 | **4.389\*a** | 2.519 |
| ACC LH | -.059 ± .119 | -.080 ± .094 | -.095 ± .106 | **5.019\*a** | -.092 ± .109 | -.088 ± .116 | -.089 ± .105 | .036 | 3.195<sup>+</sup> |
| ACC RH | -.056 ± .119 | -.068 ± .102 | -.080 ± .111 | 2.026 | -.085 ± .107 | -.089 ± .100 | -.085 ± .097 | 0.000 | 1.179 |
| Fusiform LH | .758 ± .258 | .743 ± .207 | .553 ± .120 | **65.492\*ab** | .705 ± .285 | .724 ± .294 | .579 ± .259 | **22.579\*ab** | **4.803\*** |
| Fusiform RH | .855 ± .303 | .851 ± .234 | .669 ± .229 | **49.514\*ab** | .823 ± .292 | .845 ± .301 | .694 ± .253 | **49.438\*ab** | 3.322<sup>+</sup> |

Results from the ROI analyses of distractor effects indicated strong effects of distractor type on neural activation patterns. Specifically, among all participants, significantly higher responses were observed in several regions (EROS, left DLPFC, left VLPFC, right OFC, bilateral parahippocampal gyrus, bilateral hippocampus, bilateral amygdala, and bilateral fusiform) to both alcbev and nonalcbev relative to scrambled stimuli. The effect was significantly larger for the ALC group than the NC group in the left EROS and left fusiform regions, and in four prefrontal brain areas (left DLPFC, left VLPFC, left OFC, and right OFC). Additionally, in the alcbev vs. nonalcbev contrast, the ALC group showed significantly higher responses than the NC group in the left OFC ROI. Figure 5 shows percent signal change over time for the OFC, VLPFC, fusiform, and ACC activation, to demonstrate representative activity patterns. The left OFC shows the heightened activation for alcbev for the ALC group, the fusiform and VLPFC show higher activation in both groups to both beverage types, and the ACC demonstrates how activation is lower during distractor presentation than during fixation (for both groups and for all three distractor types).

**Figure 5.**
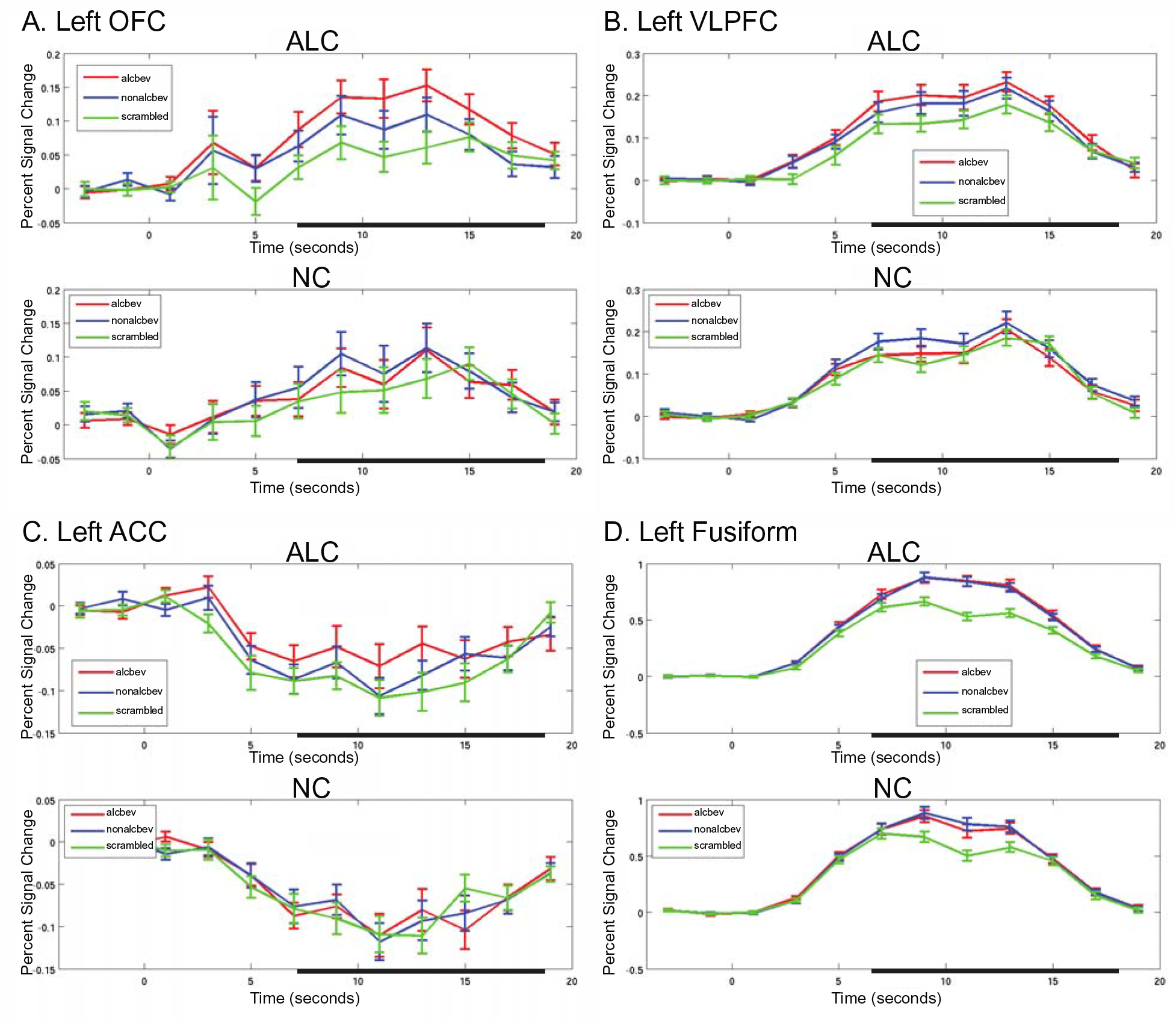
ROI - Percent signal changes in regions of interest for ALC and NC participants by distractor type. The percent signal change represents brain activity during presentation of fixation and the task stimuli. Error bars represent the standard error of the mean. Time zero was set to the onset of the encoded faces, and signal zero was set to the average signal for the three initial time points (two pre-trial and one post-trial onset). The analysis window used to examine the distractor was 6 to 16 seconds following trial onset (2 to 12 seconds after distractor onset), as indicated by the thick line on the x axis. Abbreviations: OFC = orbitofrontal cortex; VLPFC = ventrolateral prefrontal cortex; ACC = anterior cingulate cortex. The ACC (Panel C) is part of the anterior hub within the fixation-regions. The remaining areas are primarily task-regions.

Applying a Bonferroni correction accounting for all 25 tests would set a critical value for statistical significance of main effects and Distractor, Group, and Gender interactions at *p* < 0.002. Using this threshold, main effects of Distractor would remain significant in left OFC, bilateral parahippocampal gyrus, bilateral hippocampus, right amygdala, and bilateral fusiform gyrus. All main effects of Group and Gender, as well as all interaction effects were *p* > 0.002.

## 4. Discussion

### 4.1 Behavioral Responses to Probe Face and Distractor Cues

In this study, ALC and NC participants alike were able to use emotional face valence information to improve face memory, as assessed by a DMTS task (LeDoux, 1996; Dolcos *et al*., 2005; Dolcos & McCarthy, 2006). This was evidenced by better memory performance (*i.e.*, higher accuracy) on positive and negative faces, and faster reaction times to positive faces, than neutral faces. Because it has been shown that alcoholism is associated with impaired emotional perception, and specifically impaired emotional face decoding (Oscar-Berman *et al*., 1990; Philippot *et al*., 1999; Clark *et al*., 2007; Marinkovic *et al*., 2009; Hoffman *et al*., 2019; Lewis *et al*., 2019), we had postulated that normal enhancement of memory by emotional content (in this case, emotional facial expressions) would not be as strong in the ALC group. In those studies, the ALC groups were mostly or exclusively men. However, our results suggest that the ability to use emotional information to aid face memory implicitly may be relatively preserved in AUDw. A significant Group-by-Gender interaction was observed on recognition accuracy as shown in Figure A1 and Table A1. Although the Group-by-Gender interaction for reaction times was not significant (Figure A2 and Table A2), the pattern is congruent with the accuracy data. The higher accuracy of the ALCw (∼6 percentage points better than ALCm or NCw) may reflect their greater focus and sensitivity to emotional faces and resistance to distraction or underlying differences in personality or motivation (Mosher Ruiz *et al*., 2017).

We had postulated that the ALC group would show more recognition errors after alcohol distractors (relative to other distractor types), whereas the NC group would not. However, we found that regardless of the group, the distractor type did not substantially influence accuracy or reaction time. Although we also had expected performance by the ALC group to be impaired by alcoholic beverage cues, a significant interaction between group and distractor type was not evident. We did not find an attentional bias effect: The alcoholic beverage distractors (relative to nonalcoholic beverage or scrambled picture distractors) did not disproportionately decrease the number of correctly recalled faces among either the ALC or the NC groups. A review by Field and Cox (2008) suggested that the strength of the bias among ALC groups depends on drinking history. Interestingly, Loeber and colleagues (2009) found that reduced attentional biasing to alcohol cues was associated with longer durations of heavy drinking, and our sample also had long durations. Combining participants with variable drinking histories might have masked the attentional biasing effect. However, measures of neural activity may be more sensitive than behavioral measures to changes associated with long-term heavy drinking.

### 4.2 Distractor fMRI Contrasts

The fMRI contrasts revealed broadly similar patterns of brain activity among the four groups, for both fixation-regions and task-regions. In both brain networks, the beverages (alcbev and nonalcbev) elicited higher activation than the scrambled distractors. This amounts to internal replications of our present fMRI results, with four independent samples (ALCm, ALCw, NCm, and NCw) revealing the same fixation-regions, same task-regions, and with mostly the same direction of effects for task contrasts within those regions. The results reflect the existence, location, and extent of the DMN (Buckner & DiNicola, 2019; Uddin *et al*., 2019), and also indicate that beverage pictures elicit higher activation than scrambled pictures in the DMN. Moreover, our results indicate that DMN regions are sensitive to the informational content of visual stimuli. In task-regions, the occipital lobe, along with adjoining visual areas in temporal and parietal cortex, were clearly more activated by beverage stimuli than by scrambled images. The results further suggest that content processing is not solely performed by those visual regions, because activation to beverage cues was identified in middle to posterior cingulate regions, which are involved in a multitude of cognitive functions (Heilbronner & Hayden, 2016; Yeo *et al*., 2016).

### 4.3 Gender Differences

The present research provides further evidence for the importance of considering gender when exploring effects of alcoholism on the brain (Mann *et al*., 2005; Ruiz *et al*., 2013). Many factors contribute to the observed differences in function for abnormalities identified comparing ALCm with NCm vs ALCw with NCw.

Cortical group-level cluster analyses revealed significant Group-by-Gender interaction effects in 22 clusters. The general pattern of those findings indicated that ALCm had lower activation contrasts than NCm, while ALCw had higher activation contrasts than NCw. This pattern was observed primarily in contrasts between beverage and scrambled distractor conditions, and they were found in the two core medial DMN regions, as well as in visual association cortices. A similar pattern of results was found in a previous report (Sawyer *et al*., 2019) in which emotional vs. neutral image contrasts were lower in ALCm than NCm, and stronger in ALCw than NCw. The lower brain reactivity for ALCm, and higher for ALCw, highlighted gender effects, suggesting possible differences in the underlying basis for development of AUD. Of note, the results from other modalities also have indicated similar directions of the fMRI effects, with ALCw having larger reward regions than NCw, and higher fractional anisotropy than NCw, as compared to the smaller regions and lower fractional anisotropy found for ALCm than NCm (Sawyer *et al*., 2017, 2018).

The gender-divergent abnormalities in the anterior and posterior hub regions of the DMN could be reflective of other gender differences observed in conjunction with AUD. The role of these regions in internal monitoring could relate to differences in pre-existing risk factors (Ruiz & Oscar-Berman, 2015; Brighton *et al*., 2016; Mosher Ruiz *et al*., 2017), or could represent differential consequences of alcohol abuse (Merrill & Read, 2010). A similar pattern of group differences was identified in cortical regions associated with visual processing. That is, the results could represent a more fundamental impact that is not regionally-specific. In the present study, effects of both increased activation in reward regions and decreased deactivation in DMN regions in response to alcohol pictures were strongest among ALCw in particular. One reason stronger alcohol cue-specific responses were observed among ALCw could be related to gender-based differences in physiological responses to alcohol cues (Rubio *et al*., 2013). Aligned with this, larger responses to alcohol cues by female social drinkers relative to male social drinkers have been reported in superior and middle frontal gyri (Seo *et al*., 2011).

Another explanation for greater effects of alcohol cues among women than men could be related to depression (Saraceno *et al*., 2012). Symptoms of depression among non-treatment seeking heavy drinkers were reported to be correlated with increased activation in response to alcohol cue exposure in the insula, cingulate, ventral tegmentum, striatum, and thalamus (Feldstein Ewing *et al*., 2010). Because ALCw tend to experience depression and anxiety symptoms (Benishek *et al*., 1992; Schulte *et al*., 2009), we expected to see higher responses to alcohol cues among ALCw than ALCm in these regions. Indeed, in our sample, ALCw had higher Hamilton Rating Scale for Depression scores than did ALCm, although the scores for men and women were low.

### 4.4 Group Differences

In addition to significant gender interactions, we identified regions with differences between the ALC and NC groups. For the simple comparisons of the ALC group with the NC group, cluster analyses showed that the ALC group had higher activation in 2 fixation-regions and 13 task-regions (Table A5). Differences in the posterior hub were identified in regions with stronger activation to nonalcoholic beverages than to scrambled images, while the other clusters had stronger activation to alcoholic beverages. The higher contrasts observed for the ALC group indicate a processing bias toward beverage cues across fixation- and task-regions. Further, the fact that this cue-sensitivity is not isolated to a single region in the brain likely reflects a widespread divergence in emotional and cognitive activity.

### 4.5 Brain Responsivity in Fixation-Regions (Default Network)

Compared to the NC group, cluster analyses showed that the ALC group, and the ALCw in particular, had stronger contrasts in the anterior and posterior hubs, along with the temporoparietal junction. The results for ROI that include DMN regions also support the finding of contrast dampening in response to alcohol cues. Abnormal DMN functioning has been observed in other addictions and neuropsychiatric conditions (Broyd *et al*., 2009; Bednarski *et al*., 2011; Zhou *et al*., 2020). In AUD, abnormal functional connectivity among DMN regions has been reported (Chanraud *et al*., 2011). ALCw in particular had stronger contrasts for both the anterior and posterior hubs, an abnormality which could indicate a limitation in the level of detail processed, or the way in which it is integrated (Sormaz *et al*., 2018). Lower activation of the anterior hub specifically has been associated with dynamic attention allocation during task executions (Koshino *et al*., 2011), suggesting that reduced deactivation of this region during viewing of alcoholic beverage pictures among alcoholics could be associated with failure to reallocate attention back to the task when the alcohol distractors were presented.

### 4.6 Brain Responsivity in Regions of Interest

The ROI analysis of the OFC provides evidence for reward-specific processing in the ALC group. In particular, reward-specific processing refers to their higher activation from the contrast between alcbev and nonalcbev. Several studies have reported enhanced OFC activation to alcohol cues (Wrase *et al*., 2002; Myrick *et al*., 2008; Ray *et al*., 2010; Shields & Gremel, 2020), and research has established the role of the OFC in alcohol and drug addiction more generally (Goldstein & Volkow, 2002). The OFC activity may be particularly important for preoccupation and anticipation stages of the addiction cycle (Koob & Volkow, 2010). Additionally, activity in this region has been shown to correlate with subjective craving ratings of viewed alcohol cues (Myrick *et al*., 2004), and further correlated with relapse risk (Reinhard *et al*., 2015).

In many ROI, differences in activation levels among beverage distractor conditions (alcoholic and nonalcoholic) were larger relative to scrambled pictures in the ALC group than in the NC group. The higher responsivity of the ALC group to alcoholic beverages supports our hypothesis of greater attentional bias in the form of stronger alcbev vs. nonalcbev activity contrasts, but we did not expect the ALC group to have greater activation than the NC group to nonalcoholic beverages relative to scrambled cues. One explanation for this result is that many of the nonalcoholic beverages contain caffeine or sugar (*e.g.*, coffee, tea, soda), which, like alcoholic beverages, also stimulate reward-network activity (Garber & Lustig, 2011). As was suggested in an earlier meta-analysis (Field *et al*., 2009), the attentional bias for caffeine-related cues may correlate more strongly with subjective craving than for alcohol-related cues. Moreover, craving for both caffeine and alcohol utilize similar neural circuits as are used for processing alcohol reward (Kunz *et al*., 2008), as do the effects of sugar-related reward (Avena *et al*., 2008; Volkow *et al*., 2013).

The regions that responded more to beverage cues relative to the scrambled pictures were the total EROS and many of its subcomponents (DLPFC, VLPFC, OFC, parahippocampal gyrus, hippocampus, amygdala) and fusiform. In all of the regions where an interaction of Distractor type and Group was identified, the distractor effect was found to be significant among the ALC group, but not among controls. In the EROS, DLPFC, VLPFC, and OFC, the distractor effect in the ALC group was driven by greater activity during both alcoholic and nonalcoholic beverage pictures relative to scrambled pictures. In the OFC, however, the alcohol distractors elicited more activity in the ALC group than did the nonalcoholic beverages or the scrambled pictures. Thus, the strongest ROI effect specific to processing alcohol cues was observed in the OFC. Crucially, this effect was observed only in the ALC group and not in the NC group (*i.e.*, for controls, no ROI were identified where alcohol distractor pictures elicited more activity than did nonalcoholic beverages; see Table 3). The OFC is believed to play a major role in craving and reward function (Koob & Volkow, 2010).

Responses to alcohol and nonalcoholic beverage pictures in the fusiform gyrus were strong among the ALC and NC groups (Figure 5D), as expected, given the fusiform’s role in visual object recognition (Pourtois *et al*., 2009). We hypothesized further that the ALC group’s decreased BOLD signal in the fusiform gyrus in response to alcohol cues would provide evidence for stronger distractor interference with face memory maintenance. However, our results showed that any diminishment of the BOLD signal in the fusiform was far outweighed by its initially higher responses to the alcohol cues. While reductions in fusiform response across the time window of distractor analysis are apparent in the activation time course, an alcohol-specific decrement in the ALC group was not clear. The fact that BOLD responses in the fusiform ROI were consistently lower for scrambled pictures relative to alcohol and nonalcoholic beverage pictures in both the ALC and NC groups suggests that the differences in visual processing demands between conditions may have overridden any potential reduction in the BOLD signal as a result of distractor interference.

We hypothesized that a failure to inhibit distractor interference in response to alcohol cues would be associated with lower activity in VLPFC, given this region’s role in inhibition of task-irrelevant distracting stimuli (Thompson-Schill *et al*., 2002; Aron *et al*., 2004). However, our ROI results showed similarly increased activation of this region for both alcohol and nonalcoholic beverage cues relative to scrambled pictures. This finding suggests that rather than a failure of VLPFC to inhibit distracting stimuli, the higher activity in this region might result from the overriding demand of emotional and reward salience of the alcohol cues. Alternatively, the VLPFC could be involved in inhibition regardless of the distractor type employed during the DMTS delay.

### 4.7 Limitations

It is not clear to what degree the abnormalities we observed result from or predate heavy drinking. The mean abstinence period for the ALC group was 8.3 years, and since the NC group did not have an ‘abstinence period,’ we could not covary for sobriety. Still, our AUD cohort had drinking history values representative of the national population (World Health Organization, 2019), which thereby improves the generalizability of our results. Sobriety in our subject cohort points to how persistent the processing deficits in AUD populations are, and how short- and long-term abstinence may have different paths of recovery for men and women (Fama *et al*., 2020). Nonetheless, our findings illustrate how critical it is to pursue research examining gender differences regarding attentional bias towards reward-related stimuli and pathological alcohol consumption.

In conjunction with the multiple-comparison cluster correction procedures employed, the significance level we used (*p* < .05) has been shown to have higher false-positive rates than expected (Eklund *et al*., 2016). However, stricter thresholds would increase the chance of false-negative errors, and the significance level we used allows the size of the gender effects to be highlighted. Although we report cluster labels by the location of the peak voxel or vertex, the clusters reported can be understood to span multiple functional regions (Woo *et al*., 2014). That is, they are not limited to a single region, as reported by the maximal vertex or voxel.

Finally, our analyses did not include factors such as cigarette smoking, body mass index, and hormone therapy (Luhar *et al*., 2013; Oscar-Berman *et al*., 2014), which could possibly influence alcohol cue processing, reward, and DMN activity.

## 5. Conclusions

Compared to the NC group, the ALC group had stronger activation for DMN regions, and overactivated reward regions during alcohol cue distraction. This suggests that attentional capture is not limited to reward regions, but also includes DMN regions. If so, the DMN has a role in processing salient aspects of addictive substances.

The present study showed that alcohol cue distractors have powerful effects on reward-related regions of the brain, even in the absence of impaired performance when alcohol cues are employed as distracting stimuli. We also demonstrated that the increased responses in reward regions are accompanied by dampened DMN activity during the presentation of alcohol cues. Our results suggest that these effects are strongest among ALCw, and provide evidence for dimorphic patterns of responses to alcohol cues between ALCm and ALCw.

## Acknowledgements

This work was supported by funds from the US Department of Veterans Affairs Clinical Science Research and Development grant I01CX000326; National Institute on Alcohol Abuse and Alcoholism (NIAAA) of the National Institutes of Health US Department of Health and Human Services under Award Numbers R01AA007112, R01AA016624, K05AA00219, and K01AA13402; and shared instrumentation grants 1S10RR023401, 1S10RR019307, and 1S10RR023043 from the National Center for Research Resources (now National Center for Advancing Translational Sciences) at the Athinoula A. Martinos Center, Massachusetts General Hospital. Alcoholic and nonalcoholic beverage pictures were a combination of images used with permission from the Normative Appetitive Picture System (NAPS) (Stritzke et al., 2004), and other previously published works on alcohol cues (Wrase et al., 2002; Myrick et al., 2004). The authors thank Elinor Artsy, Sheeva Azma, Anne-Mette Guldberg, Zoe Gravitz, Doug Greve, Steve Lehar, Diane Merritt, Alan Poey, Elizabeth Rickenbacher, Trinity Urban, Maria Valmas, and Robert Zondervan for assistance with consultation, manuscript preparation and recruitment, assessment, preparing testing materials, and neuroimaging of the research participants. Finally, we would like to acknowledge the role of the research participants for making this study possible. The content is solely the responsibility of the authors and does not necessarily represent the official views of the National Institutes of Health, the U.S. Department of Veterans Affairs, or the United States Government.

## Appendix

**Figure A1.**
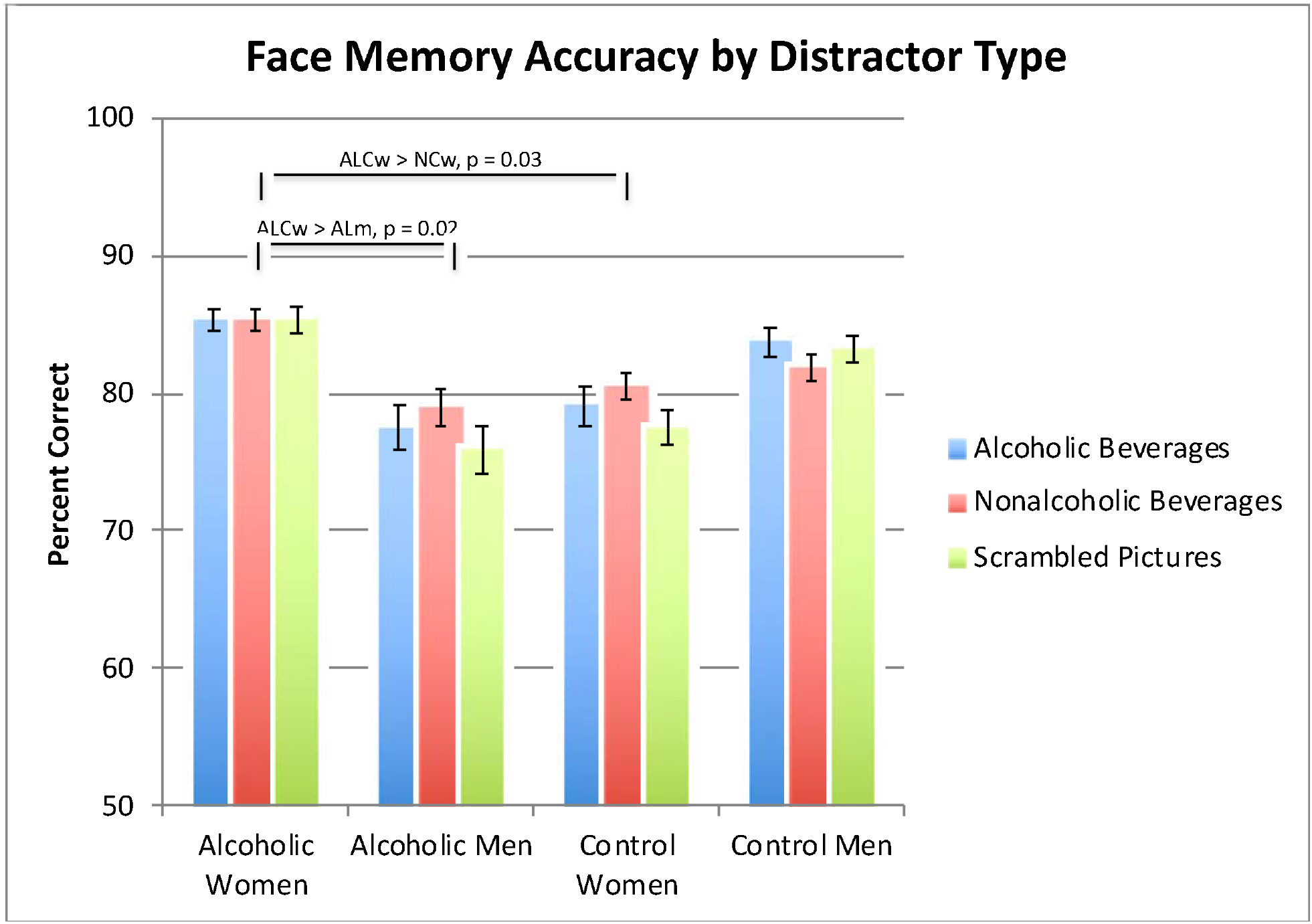
Group and gender comparisons in percent correct responses to the probe face. A significant Group-by-Gender interaction showed that face memory accuracy was significantly higher for the Alcoholic Women than the Alcoholic Men, a gender difference that was greater than the one observed for the NC group. Alcoholic Women also had higher accuracy than Control Women. (Also see Table A1.) There was no significant effect of Distractor type on performance accuracy. Error bars represent standard error of the mean.

**Figure A2.**
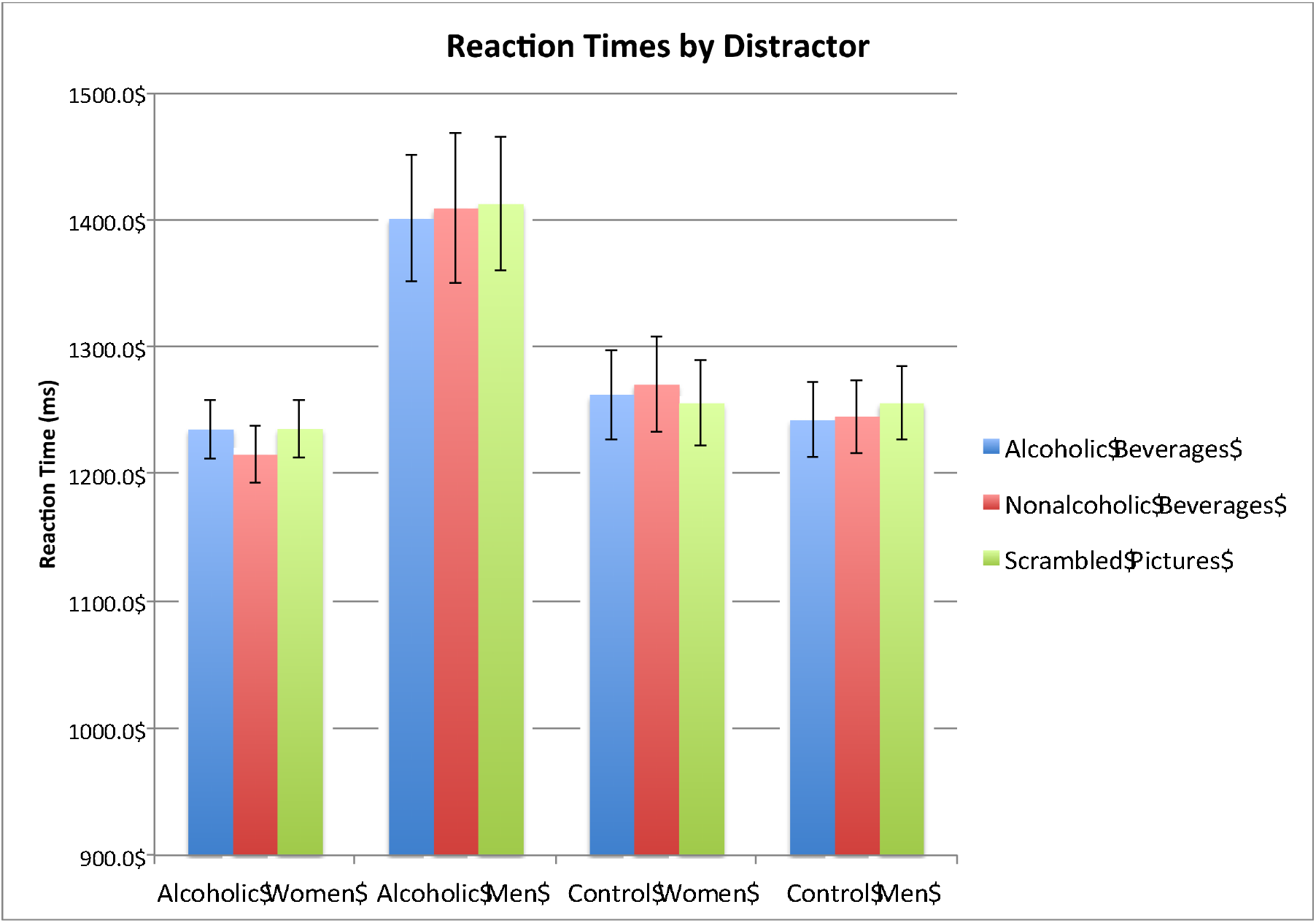
Face memory reaction times: group and gender comparisons in reaction times to the probe face as a function of the distractor stimuli. Reaction times (in milliseconds) are shown for correct trials sorted by conditions and groups. (Also see Table A2.) Participants did not vary significantly by Group or Gender on overall reaction times. There was no significant effect of Distractor type on reaction time, nor did reaction time performance by Distractor type significantly vary by Group or Gender. Error bars represent standard error of the mean.

**Figure A3.**
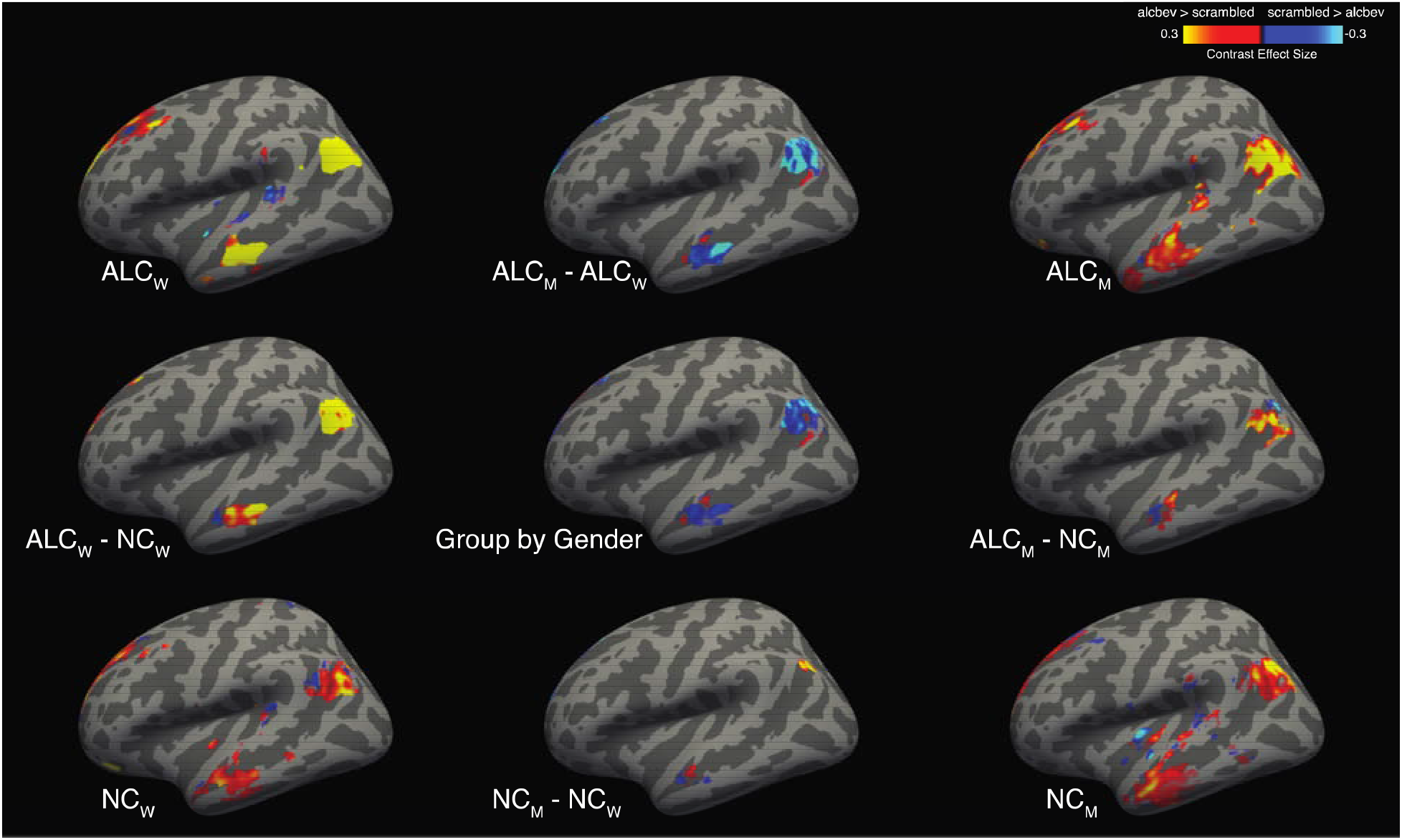
Cortical activation maps for the alcbev vs. scrambled contrast within fixation-regions (left lateral view) A significant Group-by-Gender interaction revealed several clusters (see Figure 3 and Table A4), although no clusters with significant interactions were found on the left lateral surface. Group mean contrast values (for alcbev vs. scrambled within fixation-regions) are displayed in the four brain images located in the corners of the figure, and group comparisons are indicated by minus signs. Abbreviations: ALCm = Alcoholic men; ALCw = Alcoholic women; NCm = Nonalcoholic men; NCw = Nonalcoholic women.

**Figure A4.**
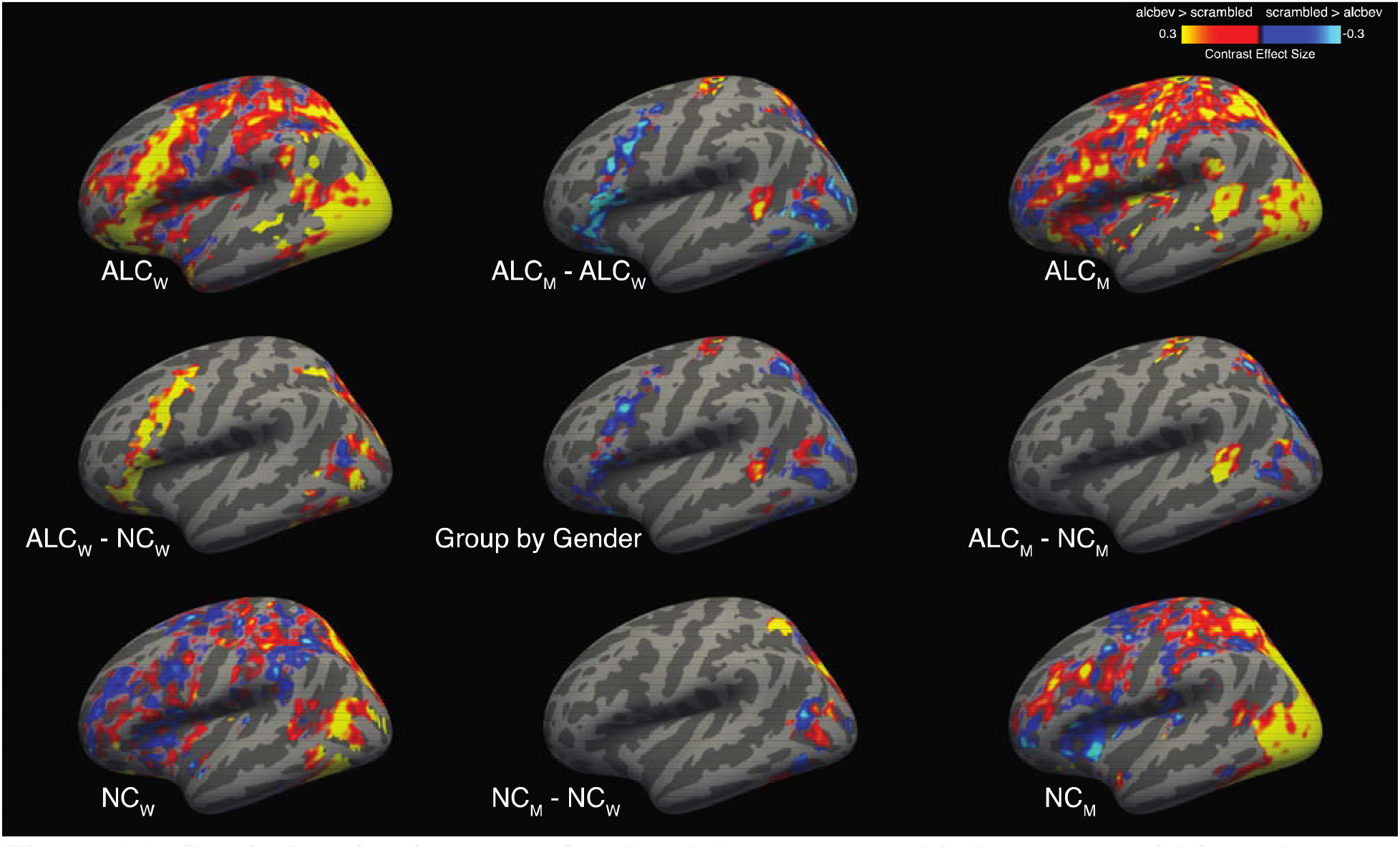
Cortical activation maps for the alcbev vs. scrambled contrast within task-regions (left lateral view) A significant Group-by-Gender interaction revealed several clusters (see Figure 4 and Table A4), although none are visible on the left lateral view. Group mean contrast values (for alcbev vs. scrambled within fixation-regions) are displayed in the four brain images located in the corners of the figure, and group comparisons are indicated by minus signs. Abbreviations: ALCm = Alcoholic men; ALCw = Alcoholic women; NCm = Nonalcoholic men; NCw = Nonalcoholic women.

**Table A1.** Behavioral task percent correct responses. Scores for accuracy are provided for overall performance, distractor type, and facial emotion. ^a^Group x Gender; ^b^Alcoholic Women > Alcoholic Men; ^c^Alcoholic Women > Control Women; ^d^Emotion main effect; ^e^Positive Faces > Neutral Faces; ^f^Negative Faces > Neutral Faces; all *p* < 0.5. Abbreviations: ALCw = Alcoholic Women; ALCm = Alcoholic Men; NCw = Nonalcoholic Control Women; NCm = Nonalcoholic Control Men.

| Percent Correct Responses Overall<sup>abc</sup> |  |  |  |  |  |  |
| --- | --- | --- | --- | --- | --- | --- |
|  | ALC | ALCw | ALCm | NC | NCw | NCm |
| *Mean* | 81.4 | 85.3 | 77.4 | 81.0 | 79.0 | 82.9 |
| *Standard Deviation* | 11.5 | 6.8 | 13.8 | 9.7 | 10.7 | 8.4 |
| *Range* | 50.0-97.5 | 66.0-96.3 | 50.0-97.5 | 54.9-96.9 | 54.9-92.6 | 71.0-96.9 |
| Percent Correct Responses by Distractor |  |  |  |  |  |  |
|  | ALC | ALCw | ALCm | NC | NCw | NCm |
| Alcoholic Beverages |  |  |  |  |  |  |
| *Mean* | 81.4 | 85.3 | 77.5 | 81.4 | 79.1 | 83.7 |
| *Standard Deviation* | 12.4 | 7.4 | 15.0 | 11.4 | 12.9 | 9.5 |
| *Range* | 45.8-98.1 | 66.7-96.3 | 45.8-98.1 | 51.9-98.1 | 51.9-98.1 | 64.8-98.1 |
| Nonalcoholic Beverages |  |  |  |  |  |  |
| *Mean* | 82.1 | 85.3 | 78.9 | 81.1 | 80.5 | 81.8 |
| *Standard Deviation* | 10.8 | 7.7 | 12.5 | 9.0 | 9.2 | 9.0 |
| *Range* | 53.7-98.1 | 68.8-98.1 | 53.7-96.3 | 57.4-94.4 | 57.4-91.7 | 64.8-94.4 |
| Scrambled Pictures |  |  |  |  |  |  |
| *Mean* | 80.6 | 85.3 | 75.9 | 80.4 | 77.5 | 83.2 |
| *Standard Deviation* | 13.6 | 8.6 | 16.1 | 11.1 | 12.1 | 9.4 |
| *Range* | 44.4-98.1 | 62.5-98.1 | 44.4-98.1 | 51.9-98.1 | 51.9-91.7 | 61.1-98.1 |
| Percent Correct Responses by Facial Emotion<sup>def</sup> |  |  |  |  |  |  |
|  | ALC | ALCw | ALCm | NC | NCw | NCm |
| Positive Faces |  |  |  |  |  |  |
| *Mean* | 82.4 | 86.8 | 78.1 | 80.7 | 79.4 | 82.0 |
| *Standard Deviation* | 12.5 | 9.0 | 14.2 | 10.4 | 11.9 | 8.7 |
| *Range* | 51.9-100.0 | 56.3-100.0 | 51.9-96.3 | 51.9-96.3 | 51.9-94.4 | 70.4-96.3 |
| Negative Faces |  |  |  |  |  |  |
| *Mean* | 81.9 | 85.9 | 77.8 | 82.9 | 80.9 | 84.9 |
| *Standard Deviation* | 12.0 | 7.2 | 14.4 | 9.5 | 11.0 | 7.5 |
| *Range* | 46.3-98.1 | 68.8-98.1 | 46.3-98.1 | 53.7-98.1 | 53.7-94.4 | 72.2-98.1 |
| Neutral Faces |  |  |  |  |  |  |
| *Mean* | 79.8 | 83.2 | 76.4 | 79.3 | 76.7 | 81.9 |
| *Standard Deviation* | 11.8 | 7.5 | 14.3 | 11.2 | 11.1 | 10.9 |
| *Range* | 51.9-98.1 | 62.5-92.6 | 51.9-98.1 | 50.0-96.3 | 50.0-92.6 | 63.0-96.3 |

**Table A2.** Reaction times. Group reaction times in milliseconds are provided for distractor type, facial emotion, and face gender. Footnotes indicate significant differences, all *p* < 0.5: ^a^Group x Gender interaction; ^b^Alcoholic Women > Alcoholic Men; ^c^Alcoholic Women > Control Women; ^d^Emotion main effect; ^e^Positive Faces > Neutral Faces; ^f^Negative Faces > Neutral Faces. Abbreviations: ALCw = Alcoholic Women; ALCm = Alcoholic Men; NCw = Nonalcoholic Control Women; NCm = Nonalcoholic Control Men.

|  | ALC<br>n = 42 | NC<br>n = 42 | ALCw<br>n = 21 | ALCm<br>n = 21 | NCw<br>n = 21 | NCm<br>n = 21 |
| --- | --- | --- | --- | --- | --- | --- |
| All Trials (ms) |  |  |  |  |  |  |
| mean | 1374.0 | 1302.9 | 1263.4 | 1484.6 | 1319.3 | 1286.5 |
| standard deviation | 418.2 | 300.0 | 213.0 | 536.1 | 337.2 | 265.0 |
| range | 896.6 - 2720.8 | 799.6 - 2164.9 | 896.6 - 1699.5 | 922.8 - 2720.8 | 931.2 - 2164.9 | 799.6 - 1812.7 |
| Correct Trials (ms) |  |  |  |  |  |  |
| mean | 1317.6 | 1254.9 | 1229.4 | 1405.9 | 1263.1 | 1246.7 |
| standard deviation | 377.0 | 288.1 | 207.9 | 481.5 | 321.1 | 258.7 |
| range | 889.7 - 2454.1 | 784.2 - 2129.2 | 889.7 - 1649.3 | 912.4 - 2454.1 | 909.0 - 2129.2 | 784.2 - 1795.1 |
| Incorrect Trials (ms) |  |  |  |  |  |  |
| mean | 1686.2 | 1564.2 | 1532.4 | 1840.0 | 1591.0 | 1537.5 |
| standard deviation | 538.2 | 328.2 | 285.2 | 680.3 | 376.8 | 278.1 |
| range | 1079.8 - 3883.9 | 1064.2 - 2480.1 | 1160.8 - 2374.8 | 1079.8 - 3883.9 | 1064.2 - 2480.1 | 1097.1 - 1939.1 |
|  | ALC | NC | ALCw | ALCm | NCw | NCm |
| Alcoholic Beverages (ms) |  |  |  |  |  |  |
| mean | 1318.3 | 1252.2 | 1235.4 | 1401.1 | 1261.7 | 1242.7 |
| standard deviation | 360.9 | 292.1 | 216.4 | 453.6 | 320.2 | 268.6 |
| range | 866.2 - 2367.9 | 765.7 - 2101.9 | 866.2 - 1578.8 | 926.1 - 2367.9 | 906.8 - 2101.9 | 765.7 - 1837.5 |
| Nonalcoholic Beverages (ms) |  |  |  |  |  |  |
| mean | 1312.4 | 1257.5 | 1215.5 | 1409.3 | 1270.6 | 1244.5 |
| standard deviation | 419.1 | 304.9 | 208.2 | 545.0 | 346.9 | 264.4 |
| range | 886.8 - 2682.0 | 779.7 - 2274.0 | 886.8 - 1686.6 | 902.8 - 2682.0 | 910.8 - 2274.0 | 779.7 - 1894.5 |
| Scrambled Pictures (ms) |  |  |  |  |  |  |
| mean | 1324.2 | 1255.9 | 1235.7 | 1412.7 | 1255.8 | 1256.0 |
| standard deviation | 378.8 | 281.8 | 213.1 | 482.0 | 306.5 | 262.5 |
| range | 888.7 - 2678.9 | 806.6 - 2004.1 | 914.9 - 1686.1 | 888.7 - 2678.9 | 882.4 - 2004.1 | 806.6 - 1825.8 |
|  | ALC | NC | ALCw | ALCm | NCw | NCm |
| Positive Faces (ms) |  |  |  |  |  |  |
| mean | 1290.5 | 1241.2 | 1227.9 | 1353.1 | 1238.4 | 1244.0 |
| standard deviation | 347.2 | 279.5 | 229.3 | 431.6 | 306.7 | 257.0 |
| range | 867.7 - 2608.6 | 789.2 - 2015.1 | 867.7 - 1617.1 | 894.3 - 2608.6 | 870.0 - 2015.1 | 789.2 - 1871.9 |
| Negative Faces (ms) |  |  |  |  |  |  |
| mean | 1303.9 | 1251.2 | 1225.7 | 1382.0 | 1264.3 | 1238.0 |
| standard deviation | 322.5 | 294.0 | 186.2 | 407.2 | 323.2 | 269.1 |
| range | 900.0 - 2218.6 | 778.9 - 2128.5 | 900.0 - 1674.6 | 918.8 - 2218.6 | 889.4 - 2128.5 | 778.9 - 1803.4 |
| Neutral Faces (ms) |  |  |  |  |  |  |
| mean | 1356.4 | 1272.4 | 1233.6 | 1479.2 | 1286.8 | 1258.0 |
| standard deviation | 492.4 | 307.0 | 235.3 | 640.3 | 346.8 | 269.4 |
| range | 857.7 - 3051.0 | 784.4 - 2226.0 | 857.7 - 1724.7 | 887.4 - 3051.0 | 875.3 - 2226.0 | 784.4 - 1706.6 |
|  | ALC | NC | ALCw | ALCm | NCw | NCm |
| Female Faces (ms) |  |  |  |  |  |  |
| mean | 1321.8 | 1272.3 | 1237.9 | 1405.7 | 1275.2 | 1269.4 |
| standard deviation | 369.4 | 288.7 | 206.9 | 471.3 | 311.5 | 271.8 |
| range | 903.8 - 2648.9 | 812.7 - 2089.0 | 903.8 - 1692.7 | 930.6 - 2648.9 | 919.7 - 2089.0 | 812.7 - 1830.5 |
| Male Faces (ms) |  |  |  |  |  |  |
| mean | 1314.0 | 1237.0 | 1220.4 | 1407.6 | 1249.7 | 1224.3 |
| standard deviation | 403.4 | 292.6 | 212.4 | 519.7 | 332.7 | 254.0 |
| range | 872.0 - 2797.2 | 756.1 - 2159.7 | 875.7 - 1605.9 | 872.0 - 2797.2 | 880.1 - 2159.7 | 756.1 - 1826.1 |

**Table A3.**
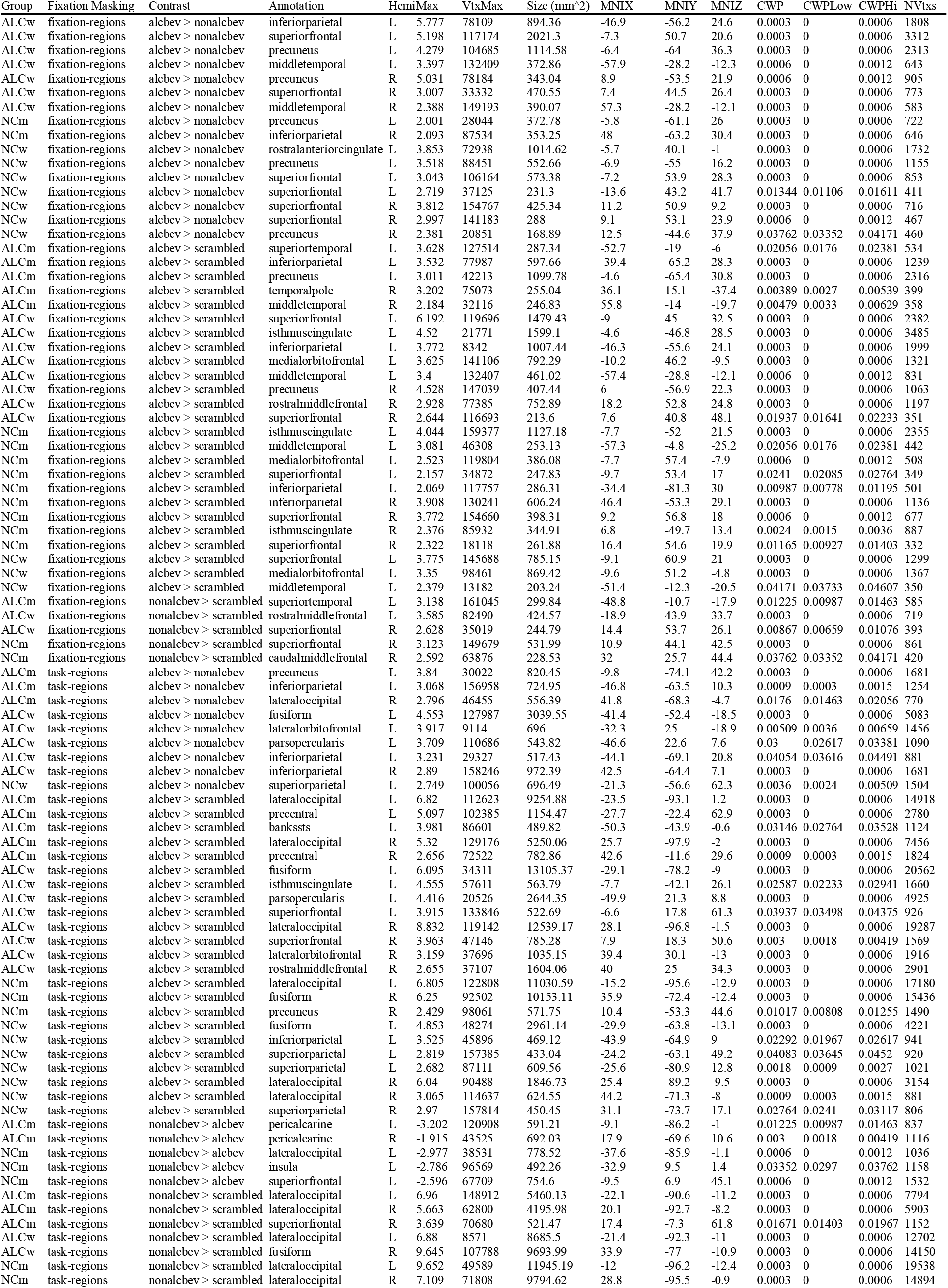
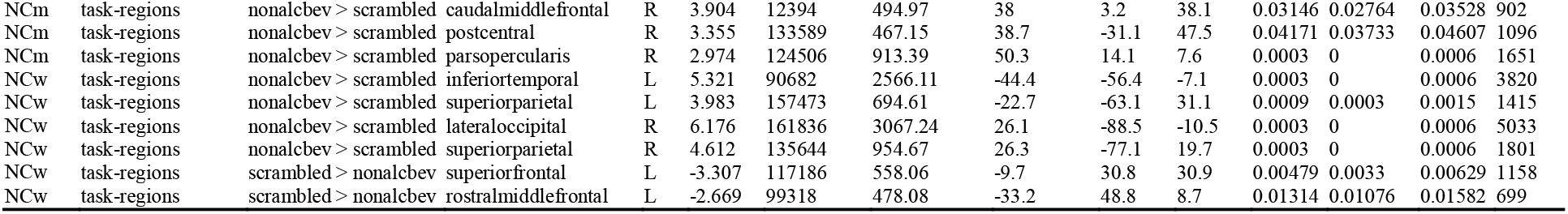
Cortical cluster characteristics for significant contrasts within each group. Annotations (from the peak voxel location in the Desikan-Killiany atlas) are shown for each of the 99 clusters with significant distractor contrasts, calculated for each group separately. The clusters reported can be understood to span multiple functional regions (Woo *et al*., 2014). That is, they are not limited to a single region, as reported by the maximal vertex or voxel. Abbreviations: Hemi = hemisphere; Max = maximum −log_10_(*p*-value) for group comparison in the cluster; VtxMax = vertex number at the maximum; Size = surface area of cluster; MNIX, MNIY, and MNIZ = Montreal Neurological Institute 305-subject template coordinates X, Y, and Z for the maximum vertex; CWP = cluster-wise *p*-value further corrected for the three spaces of left cortex, right cortex, and volume; CWPLow and CWPHi = 90% confidence intervals for CWP; NVtxs = number of vertices in the cluster; alcbev = alcoholic beverages; nonalcbev = nonalcoholic beverages; L = left hemisphere; R = right hemisphere; bankssts = banks of the superior temporal sulcus.

**Table A4.** Cortical cluster characteristics for significant Group-by-Gender interactions. Annotations (from the peak voxel location in the Desikan-Killiany atlas) are shown separately for each of the 22 clusters with significant Group-by-Gender interactions for distractor contrasts. The clusters reported can be understood to span multiple functional regions (Woo *et al*., 2014). That is, they are not limited to a single region, as reported by the maximal vertex or voxel. Abbreviations: Hemi = hemisphere; Max = maximum −log_10_(*p*-value) for group comparison in the cluster; VtxMax = vertex number at the maximum; Size = surface area of cluster; MNIX, MNIY, and MNIZ = Montreal Neurological Institute 305-subject template coordinates X, Y, and Z for the maximum vertex; CWP = cluster-wise *p*-value further corrected for the three spaces of left cortex, right cortex, and volume; CWPLow and CWPHi = 90% confidence intervals for CWP; NVtxs = number of vertices in the cluster; alcbev = alcoholic beverages; nonalcbev = nonalcoholic beverages; L = left hemisphere; R = right hemisphere. See Table 2 for a summary of the cluster information provided here.

| Fixation Masking | Contrast | Annotation | Hemi | Max | VtxMax | Size (mm<sup>2</sup>) | MNIX | MNIY | MNIZ | CWP | CWPLow | CWPHi | NVtxs |
| --- | --- | --- | --- | --- | --- | --- | --- | --- | --- | --- | --- | --- | --- |
| fixation-regions | alcbev > scrambled | superiorfrontal | L | -2.635 | 55177 | 313.89 | -7.9 | 46.2 | 22.9 | 0.01136 | 0.00897 | 0.01374 | 461 |
| fixation-regions | alcbev > scrambled | superiorfrontal | R | -3.304 | 146084 | 140.84 | 16.5 | 53.5 | 24.4 | 0.0021 | 0.0012 | 0.003 | 201 |
| fixation-regions | alcbev > scrambled | superiorfrontal | R | -2.453 | 134468 | 116.29 | 12 | 47.4 | 13.5 | 0.01017 | 0.00808 | 0.01255 | 184 |
| fixation-regions | alcbev > scrambled | precuneus | R | -1.444 | 122115 | 162.46 | 9.6 | -54.5 | 24.9 | 0.0003 | 0 | 0.0006 | 418 |
| fixation-regions | nonalcbev > scrambled | superiorfrontal | R | -3.422 | 59361 | 133.19 | 7.6 | 44.3 | 38.6 | 0.0003 | 0 | 0.0006 | 242 |
| fixation-regions | nonalcbev > scrambled | superiorfrontal | R | -3.143 | 8241 | 206.58 | 16 | 53.9 | 24.6 | 0.0003 | 0 | 0.0006 | 313 |
| fixation-regions | nonalcbev > scrambled | caudalmiddlefrontal | R | -2.747 | 29029 | 155.88 | 27.8 | 19.3 | 42.7 | 0.0003 | 0 | 0.0006 | 297 |
| task-regions | alcbev > scrambled | superiorparietal | L | -1.996 | 64642 | 333.28 | -22.2 | -84.9 | 23.6 | 0.00927 | 0.00718 | 0.01136 | 479 |
| task-regions | alcbev > scrambled | fusiform | R | -4.208 | 114347 | 1011.41 | 31.8 | -74.4 | -11.2 | 0.0003 | 0 | 0.0006 | 1562 |
| task-regions | alcbev > scrambled | superiorparietal | R | -2.983 | 157893 | 409.83 | 21.3 | -83.2 | 20.7 | 0.0012 | 0.0006 | 0.0021 | 627 |
| task-regions | alcbev > scrambled | lingual | R | -2.804 | 79452 | 375.68 | 6.9 | -68.3 | 2.4 | 0.0027 | 0.0015 | 0.00389 | 473 |
| task-regions | alcbev > scrambled | superiorparietal | R | -2.679 | 93240 | 261.8 | 24.9 | -60.1 | 48 | 0.03264 | 0.02882 | 0.03645 | 553 |
| task-regions | nonalcbev > scrambled | superiorparietal | L | -2.534 | 120369 | 478.5 | -17 | -88.7 | 20.5 | 0.0003 | 0 | 0.0006 | 709 |
| task-regions | nonalcbev > scrambled | lateraloccipital | R | -3.465 | 132542 | 658.32 | 29.9 | -85.6 | -12.5 | 0.0003 | 0 | 0.0006 | 985 |
| task-regions | nonalcbev > scrambled | superiorparietal | R | -2.61 | 130192 | 242.3 | 19.9 | -79.4 | 43 | 0.01908 | 0.01611 | 0.02204 | 404 |
| task-regions | nonalcbev > scrambled | lateraloccipital | R | -2.437 | 61711 | 238.46 | 21.3 | -87.7 | 18 | 0.02115 | 0.01789 | 0.0244 | 346 |
| task-regions | scrambled > nonalcbev | superiorfrontal | L | -3.795 | 4781 | 140.18 | -13.4 | 32.7 | 24.3 | 0.00718 | 0.00539 | 0.00897 | 238 |
| task-regions | scrambled > nonalcbev | rostralmiddlefrontal | L | -3.026 | 116230 | 173.7 | -31.1 | 48.1 | 7.1 | 0.0012 | 0.0006 | 0.0021 | 258 |
| task-regions | scrambled > nonalcbev | superiorfrontal | L | -1.97 | 153483 | 148 | -12 | 18.5 | 36.9 | 0.00449 | 0.003 | 0.00599 | 327 |
| task-regions | scrambled > nonalcbev | rostralmiddlefrontal | L | -1.861 | 36306 | 136.94 | -36.8 | 31.1 | 29.6 | 0.00897 | 0.00688 | 0.01106 | 200 |
| task-regions | nonalcbev > alcbev | superiorfrontal | L | 3.14 | 140803 | 102.82 | -8.9 | 30.4 | 32 | 0.04287 | 0.0385 | 0.04724 | 180 |
| task-regions | nonalcbev > alcbev | superiorfrontal | L | 2.283 | 110024 | 120.81 | -12.3 | 17.2 | 36.8 | 0.01403 | 0.01136 | 0.01671 | 290 |

**Table A5.**
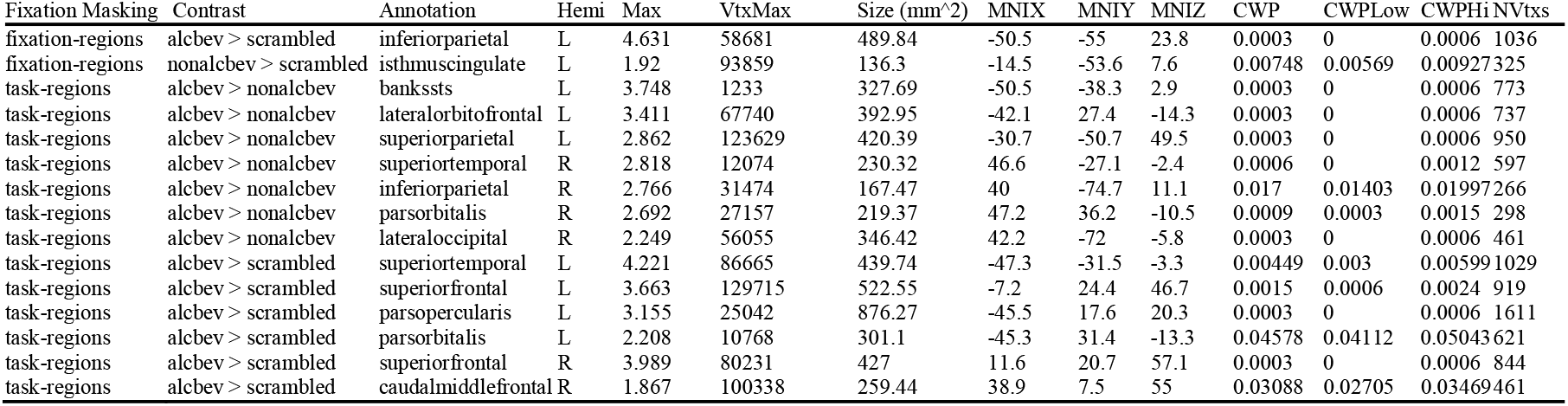
Cortical cluster characteristics for significant comparisons between ALC and NC groups. The activation levels for all contrasts were significantly greater for the ALC group than for the NC group. Annotations (from the peak voxel location in the Desikan-Killiany atlas) are shown separately for each of the 15 clusters with significant group comparisons for distractor contrasts. The clusters reported can be understood to span multiple functional regions (Woo *et al*., 2014). That is, they are not limited to a single region, as reported by the maximal vertex or voxel. Abbreviations: Hemi = hemisphere; Max = maximum −log_10_(*p*-value) for group comparison in the cluster; VtxMax = vertex number at the maximum; Size = surface area of cluster; MNIX, MNIY, and MNIZ = Montreal Neurological Institute 305-subject template coordinates X, Y, and Z for the maximum vertex; CWP = cluster-wise *p*-value further corrected for the three spaces of left cortex, right cortex, and volume; CWPLow and CWPHi = 90% confidence intervals for CWP; NVtxs = number of vertices in the cluster; alcbev = alcoholic beverages; nonalcbev = nonalcoholic beverages; L = left hemisphere; R = right hemisphere; bankssts = banks of the superior temporal sulcus.

## References

Alba-Ferrara, L., Müller-Oehring, E.M., Sullivan, E.V., Pfefferbaum, A., & Schulte, T. (2016) Brain responses to emotional salience and reward in alcohol use disorder. Brain Imaging Behav., 10, 136– 146.

Aron, A.R., Robbins, T.W., & Poldrack, R.A. (2004) Inhibition and the right inferior frontal cortex. Trends Cogn. Sci., 8, 170–177.

Avena, N.M., Rada, P., & Hoebel, B.G. (2008) Evidence for sugar addiction: behavioral and neurochemical effects of intermittent, excessive sugar intake. Neurosci. Biobehav. Rev., 32, 20–39.

Beaty, R.E., Benedek, M., Silvia, P.J., & Schacter, D.L. (2016) Creative Cognition and Brain Network Dynamics. Trends Cogn. Sci., 20, 87–95.

Becker, J.B., McClellan, M.L., & Reed, B.G. (2017) Sex differences, gender and addiction. J. Neurosci. Res., 95, 136–147.

Bednarski, S.R., Zhang, S., Hong, K.-I., Sinha, R., Rounsaville, B.J., & Li, C.-S.R. (2011) Deficits in default mode network activity preceding error in cocaine dependent individuals. Drug Alcohol Depend., 119, e51–e57.

Benishek, L.A., Bieschke, K.J., Stöffelmayr, B.E., Mavis, B.E., & Humphreys, K.A. (1992) Gender differences in depression and anxiety among alcoholics. J. Subst. Abuse, 4, 235–245.

Bordnick, P.S., Traylor, A., Copp, H.L., Graap, K.M., Carter, B., Ferrer, M., & Walton, A.P. (2008) Assessing reactivity to virtual reality alcohol based cues. Addict. Behav., 33, 743–756.

Briggs, G.G. & Nebes, R.D. (1975) Patterns of hand preference in a student population. Cortex, 11, 230– 238.

Brighton, R., Moxham, L., & Traynor, V. (2016) Women and Alcohol Use Disorders: Factors That Lead to Harm. J. Addict. Nurs., 27, 205–213.

Broyd, S.J., Demanuele, C., Debener, S., Helps, S.K., James, C.J., & Sonuga-Barke, E.J.S. (2009) Default-mode brain dysfunction in mental disorders: a systematic review. Neurosci. Biobehav. Rev., 33, 279–296.

Buckner, R.L. & DiNicola, L.M. (2019) The brain’s default network: updated anatomy, physiology and evolving insights. Nat. Rev. Neurosci., 20, 593–608.

Cahalan, D., Cisin, I.H., & Crossley, H.M. (1969) American drinking practices: A national study of drinking behavior and attitudes. Monographs of the Rutgers Center of Alcohol Studies, 6, 260.

Carter, B.L. & Tiffany, S.T. (1999) Meta-analysis of cue-reactivity in addiction research. Addiction, 94, 327–340.

Chanraud, S., Pitel, A.-L., Pfefferbaum, A., & Sullivan, E.V. (2011) Disruption of functional connectivity of the default-mode network in alcoholism. Cereb. Cortex, 21, 2272–2281.

Clapp, W.C., Rubens, M.T., & Gazzaley, A. (2010) Mechanisms of working memory disruption by external interference. Cereb. Cortex, 20, 859–872.

Clark, U.S., Oscar-Berman, M., Shagrin, B., & Pencina, M. (2007) Alcoholism and judgments of affective stimuli. Neuropsychology, 21, 346–362.

Dale, A.M. (1999) Optimal experimental design for event-related fMRI. Hum. Brain Mapp., 8, 109–114.

Dale, A.M., Fischl, B., & Sereno, M.I. (1999) Cortical surface-based analysis. I. Segmentation and surface reconstruction. Neuroimage, 9, 179–194.

Desikan, R.S., Ségonne, F., Fischl, B., Quinn, B.T., Dickerson, B.C., Blacker, D., Buckner, R.L., Dale, A.M., Maguire, R.P., Hyman, B.T., Albert, M.S., & Killiany, R.J. (2006) An automated labeling system for subdividing the human cerebral cortex on MRI scans into gyral based regions of interest. Neuroimage, 31, 968–980.

Destrieux, C., Fischl, B., Dale, A., & Halgren, E. (2010) Automatic parcellation of human cortical gyri and sulci using standard anatomical nomenclature. Neuroimage, 53, 1–15.

Dolcos, F., LaBar, K.S., & Cabeza, R. (2005) Remembering one year later: role of the amygdala and the medial temporal lobe memory system in retrieving emotional memories. Proc. Natl. Acad. Sci. U. S. A., 102, 2626–2631.

Dolcos, F. & McCarthy, G. (2006) Brain systems mediating cognitive interference by emotional distraction. J. Neurosci., 26, 2072–2079.

Eklund, A., Nichols, T.E., & Knutsson, H. (2016) Cluster failure: Why fMRI inferences for spatial extent have inflated false-positive rates. Proc. Natl. Acad. Sci. U. S. A., 113, 7900–7905.

Fama, R., Le Berre, A.-P., & Sullivan, E.V. (2020) Alcohol’s Unique Effects on Cognition in Women: A 2020 (Re)view to Envision Future Research and Treatment. Alcohol Res., 40, 03.

Feldstein Ewing, S.W., Filbey, F.M., Chandler, L.D., & Hutchison, K.E. (2010) Exploring the relationship between depressive and anxiety symptoms and neuronal response to alcohol cues. Alcohol. Clin. Exp. Res., 34, 396–403.

Field, M. & Cox, W. (2008) Attentional bias in addictive behaviors: A review of its development, causes, and consequences. Drug and Alcohol Dependence, 97, 1–20.

Field, M. & Eastwood, B. (2005) Experimental manipulation of attentional bias increases the motivation to drink alcohol. Psychopharmacology, 183, 350–357.

Field, M., Marhe, R., & Franken, I.H.A. (2014) The clinical relevance of attentional bias in substance use disorders. CNS Spectr., 19, 225–230.

Field, M., Mogg, K., Zetteler, J., & Bradley, B.P. (2004) Attentional biases for alcohol cues in heavy and light social drinkers: the roles of initial orienting and maintained attention. Psychopharmacology, 176, 88–93.

Field, M., Munafò, M.R., & Franken, I.H.A. (2009) A meta-analytic investigation of the relationship between attentional bias and subjective craving in substance abuse. Psychol. Bull., 135, 589–607.

Fischl, B., Salat, D.H., Busa, E., Albert, M., Dieterich, M., Haselgrove, C., van der Kouwe, A., Killiany, R., Kennedy, D., Klaveness, S., Montillo, A., Makris, N., Rosen, B., & Dale, A.M. (2002) Whole brain segmentation: automated labeling of neuroanatomical structures in the human brain. Neuron, 33, 341–355.

Fischl, B., Sereno, M.I., Tootell, R.B.H., & Dale, A.M. (1999) High-resolution intersubject averaging and a coordinate system for the cortical surface. Human Brain Mapping, 8, 272–284.

Flannery, B.A., Volpicelli, J.R., & Pettinati, H.M. (1999) Psychometric properties of the Penn Alcohol Craving Scale. Alcohol. Clin. Exp. Res., 23, 1289–1295.

Franken, I.H.A. (2003) Drug craving and addiction: integrating psychological and neuropsychopharmacological approaches. Prog. Neuropsychopharmacol. Biol. Psychiatry, 27, 563– 579.

Fryer, S.L., Jorgensen, K.W., Yetter, E.J., Daurignac, E.C., Watson, T.D., Shanbhag, H., Krystal, J.H., & Mathalon, D.H. (2013) Differential brain response to alcohol cue distractors across stages of alcohol dependence. Biol. Psychol., 92, 282–291.

Garber, A.K. & Lustig, R.H. (2011) Is fast food addictive? Curr. Drug Abuse Rev., 4, 146–162.

George, M.S., Anton, R.F., Bloomer, C., Teneback, C., Drobes, D.J., Lorberbaum, J.P., Nahas, Z., & Vincent, D.J. (2001) Activation of prefrontal cortex and anterior thalamus in alcoholic subjects on exposure to alcohol-specific cues. Arch. Gen. Psychiatry, 58, 345–352.

Goldstein, R.Z. & Volkow, N.D. (2002) Drug addiction and its underlying neurobiological basis: neuroimaging evidence for the involvement of the frontal cortex. Am. J. Psychiatry, 159, 1642–1652.

Goldstein, R.Z. & Volkow, N.D. (2011) Dysfunction of the prefrontal cortex in addiction: neuroimaging findings and clinical implications. Nature Reviews Neuroscience, 12, 652–669.

Greve, D.N. & Fischl, B. (2009) Accurate and robust brain image alignment using boundary-based registration. Neuroimage, 48, 63–72.

Grüsser, S.M., Heinz, A., & Flor, H. (2000) Standardized stimuli to assess drug craving and drug memory in addicts. J. Neural Transm., 107, 715–720.

Hamilton, M. (1960) A rating scale for depression. J. Neurol. Neurosurg. Psychiatry, 23, 56–62.

Heilbronner, S.R. & Hayden, B.Y. (2016) Dorsal anterior cingulate cortex: A bottom-up view. Annu. Rev. Neurosci., 39, 149–170.

Heinz, A., Wrase, J., Kahnt, T., Beck, A., Bromand, Z., Grüsser, S.M., Kienast, T., Smolka, M.N., Flor, H., & Mann, K. (2007) Brain activation elicited by affectively positive stimuli is associated with a lower risk of relapse in detoxified alcoholic subjects. Alcohol. Clin. Exp. Res., 31, 1138–1147.

Hoffman, L.A., Lewis, B., & Nixon, S.J. (2019) Neurophysiological and interpersonal correlates of emotional face processing in Alcohol Use Disorder. Alcohol. Clin. Exp. Res., 43.

Holdnack, J.A. & Drozdick, L.W. (2010) CHAPTER 9 - Using WAIS-IV with WMS-IV. In Weiss, L.G., Saklofske, D.H., Coalson, D.L., & Raiford, S.E. (eds), WAIS-IV Clinical Use and Interpretation. Academic Press, San Diego, pp. 237–283.

Jha, A.P., Fabian, S.A., & Aguirre, G.K. (2004) The role of prefrontal cortex in resolving distractor interference. Cogn. Affect. Behav. Neurosci., 4, 517–527.

Kaag, A.M., Wiers, R.W., de Vries, T.J., Pattij, T., & Goudriaan, A.E. (2019) Striatal alcohol cue-reactivity is stronger in male than female problem drinkers. Eur. J. Neurosci., 50, 2264–2273.

Koob, G.F. & Volkow, N.D. (2010) Neurocircuitry of addiction. Neuropsychopharmacology, 35, 217– 238.

Koshino, H., Minamoto, T., Ikeda, T., Osaka, M., Otsuka, Y., & Osaka, N. (2011) Anterior medial prefrontal cortex exhibits activation during task preparation but deactivation during task execution. PLoS One, 6, e22909.

Kunz, S., Beblo, T., Driessen, M., & Woermann, F. (2008) fMRI of alcohol craving after individual cues: a follow-up case report. Neurocase, 14, 343–346.

LeDoux, J.E. (1996) The Emotional Brain: The Mysterious Underpinnings of Emotional Life. Simon & Schuster.

Lewis, B., Price, J.L., Garcia, C.C., & Nixon, S.J. (2019) Emotional Face Processing among Treatment-Seeking Individuals with Alcohol Use Disorders: Investigating Sex Differences and Relationships with Interpersonal Functioning. Alcohol Alcohol, 54, 361–369.

Loeber, S., Vollstädt-Klein, S., von der Goltz, C., Flor, H., Mann, K., & Kiefer, F. (2009) Attentional bias in alcohol-dependent patients: the role of chronicity and executive functioning. Addict. Biol., 14, 194–203.

Lubman, D.I. (2007) Addiction neuroscience and its relevance to clinical practice. Drug Alcohol Rev., 26, 1–2.

Luhar, R.B., Sawyer, K.S., Gravitz, Z., Ruiz, S.M., & Oscar-Berman, M. (2013) Brain volumes and neuropsychological performance are related to current smoking and alcoholism history. Neuropsychiatr. Dis. Treat., 9, 1767–1784.

Makris, N., Oscar-Berman, M., Jaffin, S.K., Hodge, S.M., Kennedy, D.N., Caviness, V.S., Marinkovic, K., Breiter, H.C., Gasic, G.P., & Harris, G.J. (2008) Decreased volume of the brain reward system in alcoholism. Biological Psychiatry, 64, 192–202.

Mann, K., Ackermann, K., Croissant, B., Mundle, G., Nakovics, H., & Diehl, A. (2005) Neuroimaging of gender differences in alcohol dependence: Are women more vulnerable? Alcoholism: Clinical & Experimental Research, 29, 896–901.

Margulies, D.S., Ghosh, S.S., Goulas, A., Falkiewicz, M., Huntenburg, J.M., Langs, G., Bezgin, G., Eickhoff, S.B., Castellanos, F.X., Petrides, M., Jefferies, E., & Smallwood, J. (2016) Situating the default-mode network along a principal gradient of macroscale cortical organization. Proc. Natl. Acad. Sci. U. S. A., 113, 12574–12579.

Marinkovic, K., Oscar-Berman, M., Urban, T., O’Reilly, C.E., Howard, J.A., Sawyer, K., & Harris, G.J. (2009) Alcoholism and dampened temporal limbic activation to emotional faces. Alcohol. Clin. Exp. Res., 33, 1880–1892.

Menon, V. (2011) Large-scale brain networks and psychopathology: a unifying triple network model. Trends Cogn. Sci., 15, 483–506.

Merrill, J.E. & Read, J.P. (2010) Motivational pathways to unique types of alcohol consequences. Psychol. Addict. Behav., 24, 705–711.

Mosher Ruiz, S., Oscar-Berman, M., Kemppainen, M.I., Valmas, M.M., & Sawyer, K.S. (2017) Associations between personality and drinking motives among abstinent adult alcoholic men and women. Alcohol Alcohol, 52, 496–505.

Myrick, H., Anton, R.F., Li, X., Henderson, S., Drobes, D., Voronin, K., & George, M.S. (2004) Differential brain activity in alcoholics and social drinkers to alcohol cues: relationship to craving. Neuropsychopharmacology, 29, 393–402.

Myrick, H., Anton, R.F., Li, X., Henderson, S., Randall, P.K., & Voronin, K. (2008) Effect of naltrexone and ondansetron on alcohol cue-induced activation of the ventral striatum in alcohol-dependent people. Arch. Gen. Psychiatry, 65, 466–475.

Oscar-Berman, M., Hancock, M., Mildworf, B., Hutner, N., & Weber, D.A. (1990) Emotional perception and memory in alcoholism and aging. Alcohol. Clin. Exp. Res., 14, 383–393.

Oscar-Berman, M. & Maleki, N. (2019) Alcohol Dementia, Wernicke’s Encephalopathy, and Korsakoff’s Syndrome. In Michael L. Alosco and Robert A. Stern (ed), The Oxford Handbook of Adult Cognitive Disorders. Oxford University Press, pp. 743–758.

Oscar-Berman, M., Ruiz, S.M., Marinkovic, K., Valmas, M.M., Harris, G.J., & Sawyer, K.S. (2019) Brain responsivity to emotional faces differs in alcoholic men and women. bioRxiv, 496166.

Oscar-Berman, M., Valmas, M.M., Sawyer, K.S., Ruiz, S.M., Luhar, R.B., & Gravitz, Z.R. (2014) Profiles of impaired, spared, and recovered neuropsychologic processes in alcoholism. Handb. Clin. Neurol., 125, 183–210.

Philippot, P., Kornreich, C., Blairy, S., Baert, I., Den Dulk, A., Le Bon, O., Streel, E., Hess, U., Pelc, I., & Verbanck, P. (1999) Alcoholics’ deficits in the decoding of emotional facial expression. Alcohol. Clin. Exp. Res., 23, 1031–1038.

Pourtois, G., Schwartz, S., Spiridon, M., Martuzzi, R., & Vuilleumier, P. (2009) Object representations for multiple visual categories overlap in lateral occipital and medial fusiform cortex. Cereb. Cortex, 19, 1806–1819.

Ray, S., Hanson, C., Hanson, S.J., & Bates, M.E. (2010) fMRI BOLD response in high-risk college students (Part 1): during exposure to alcohol, marijuana, polydrug and emotional picture cues. Alcohol Alcohol, 45, 437–443.

Reinhard, I., Leménager, T., Fauth-Bühler, M., Hermann, D., Hoffmann, S., Heinz, A., Kiefer, F., Smolka, M.N., Wellek, S., Mann, K., & Vollstädt-Klein, S. (2015) A comparison of region-of-interest measures for extracting whole brain data using survival analysis in alcoholism as an example. J. Neurosci. Methods, 242, 58–64.

Rivas-Grajales, A.M., Sawyer, K.S., Karmacharya, S., Papadimitriou, G., Camprodon, J.A., Harris, G.J., Kubicki, M., Oscar-Berman, M., & Makris, N. (2018) Sexually dimorphic structural abnormalities in major connections of the medial forebrain bundle in alcoholism. NeuroImage: Clinical, 19, 98–105.

Robins, L.N., Cottler, L.B., Bucholz, K.K., Compton, W.M., North, C.S., & Rourke, K. (2000) Computerized Diagnostic Interview Schedule for the DSM-IV (C DIS-IV).

Rubio, G., Martínez-Gras, I., Ponce, G., Quinto, R., Jurado, R., & Jiménez-Arriero, M.Á. (2013) [Integration of self-guidance groups for relatives in a public program of alcoholism treatment]. Adicciones, 25, 37–44.

Ruiz, S.M. & Oscar-Berman, M. (2015) Gender and alcohol abuse: history and sociology. In Martin, S.C. (ed), Gender and Alcohol Abuse: History and Sociology, The SAGE Encyclopedia of Alcohol: Social, Cultural, and Historical Perspectives. Sage Publications Los Angeles, pp. 586–591.

Ruiz, S.M., Oscar-Berman, M., Sawyer, K.S., Valmas, M.M., Urban, T., & Harris, G.J. (2013) Drinking history associations with regional white matter volumes in alcoholic men and women. Alcohol. Clin. Exp. Res., 37, 110–122.

Ryan, F. (2002) Attentional bias and alcohol dependence: a controlled study using the modified stroop paradigm. Addict. Behav., 27, 471–482.

Saraceno, L., Heron, J., Munafò, M., Craddock, N., & van den Bree, M.B.M. (2012) The relationship between childhood depressive symptoms and problem alcohol use in early adolescence: findings from a large longitudinal population-based study. Addiction, 107, 567–577.

Sawyer, K.S., Adra, N., Salz, D.M., Kemppainen, M.I., Ruiz, S.M., Harris, G.J., & Oscar-Berman, M. (2020) Hippocampal subfield volumes in abstinent men and women with a history of alcohol use disorder. PLoS One, 15, e0236641.

Sawyer, K.S., Maleki, N., Papadimitriou, G., Makris, N., Oscar-Berman, M., & Harris, G.J. (2018) Cerebral white matter sex dimorphism in alcoholism: a diffusion tensor imaging study. Neuropsychopharmacology, 43, 1876–1883.

Sawyer, K.S., Maleki, N., Urban, T., Marinkovic, K., Karson, S., Ruiz, S.M., Harris, G.J., & Oscar-Berman, M. (2019) Alcoholism gender differences in brain responsivity to emotional stimuli. Elife, 8.

Sawyer, K.S., Oscar-Berman, M., Barthelemy, O.J., Papadimitriou, G.M., Harris, G.J., & Makris, N. (2017) Gender dimorphism of brain reward system volumes in alcoholism. Psychiatry Res Neuroimaging, 263, 15–25.

Sawyer, K.S., Poey, A., Ruiz, S.M., Marinkovic, K., & Oscar-Berman, M. (2015) Measures of skin conductance and heart rate in alcoholic men and women during memory performance. PeerJ, 3, e941.

Schacht, J.P., Anton, R.F., & Myrick, H. (2013) Functional neuroimaging studies of alcohol cue reactivity: a quantitative meta-analysis and systematic review: Alcohol cue imaging. Addict. Biol., 18, 121–133.

Schneider, F., Habel, U., Wagner, M., Franke, P., Salloum, J.B., Shah, N.J., Toni, I., Sulzbach, C., Hönig, K., Maier, W., Gaebel, W., & Zilles, K. (2001) Subcortical correlates of craving in recently abstinent alcoholic patients. Am. J. Psychiatry, 158, 1075–1083.

Schulte, M.T., Ramo, D., & Brown, S.A. (2009) Gender differences in factors influencing alcohol use and drinking progression among adolescents. Clin. Psychol. Rev., 29, 535–547.

Ségonne, F., Dale, A.M., Busa, E., Glessner, M., Salat, D., Hahn, H.K., & Fischl, B. (2004) A hybrid approach to the skull stripping problem in MRI. NeuroImage, 22, 1060–1075.

Seitz, J., Sawyer, K.S., Papadimitriou, G., Oscar-Berman, M., Ng, I., Kubicki, A., Mouradian, P., Ruiz, S.M., Kubicki, M., Harris, G.J., & Makris, N. (2017) Alcoholism and sexual dimorphism in the middle longitudinal fascicle: a pilot study. Brain Imaging Behav., 11, 1006–1017.

Seo, D., Jia, Z., Lacadie, C.M., Tsou, K.A., Bergquist, K., & Sinha, R. (2011) Sex differences in neural responses to stress and alcohol context cues. Human Brain Mapping, 32, 1998–2013.

Sharma, D., Albery, I.P., & Cook, C. (2001) Selective attentional bias to alcohol related stimuli in problem drinkers and non-problem drinkers. Addiction, 96, 285–295.

Shields, C.N. & Gremel, C.M. (2020) Review of orbitofrontal cortex in alcohol dependence: A disrupted cognitive map? Alcohol. Clin. Exp. Res., 44, 1952–1964.

Sinha, R., Fox, H.C., Hong, K.A., Bergquist, K., Bhagwagar, Z., & Siedlarz, K.M. (2009) Enhanced negative emotion and alcohol craving, and altered physiological responses following stress and cue exposure in alcohol dependent individuals. Neuropsychopharmacology, 34, 1198–1208.

Sled, J.G., Zijdenbos, A.P., & Evans, A.C. (1998) A nonparametric method for automatic correction of intensity nonuniformity in MRI data. IEEE Transactions on Medical Imaging, 17, 87–97.

Sormaz, M., Murphy, C., Wang, H.-T., Hymers, M., Karapanagiotidis, T., Poerio, G., Margulies, D.S., Jefferies, E., & Smallwood, J. (2018) Default mode network can support the level of detail in experience during active task states. Proc. Natl. Acad. Sci. U. S. A., 115, 9318–9323.

Stritzke, W.G.K., Breiner, M.J., Curtin, J.J., & Lang, A.R. (2004) Assessment of substance cue reactivity: advances in reliability, specificity, and validity. Psychol. Addict. Behav., 18, 148–159.

Talairach, J. & Tournoux, P. (1988) Co-Planar Stereotaxic Atlas of the Human Brain: 3-Dimensional Proportional System : An Approach to Cerebral Imaging. G. Thieme, Stuttgart.

Tapert, S.F., Cheung, E.H., Brown, G.G., Frank, L.R., Paulus, M.P., Schweinsburg, A.D., Meloy, M.J., & Brown, S.A. (2003) Neural response to alcohol stimuli in adolescents with alcohol use disorder. Arch. Gen. Psychiatry, 60, 727–735.

Thesen, S., Heid, O., Mueller, E., & Schad, L.R. (2000) Prospective acquisition correction for head motion with image-based tracking for real-time fMRI. Magn. Reson. Med., 44, 457–465.

Thompson-Schill, S.L., Jonides, J., Marshuetz, C., Smith, E.E., D’Esposito, M., Kan, I.P., Knight, R.T., & Swick, D. (2002) Effects of frontal lobe damage on interference effects in working memory. Cogn. Affect. Behav. Neurosci., 2, 109–120.

Tops, M., Boksem, M.A.S., Quirin, M., IJzerman, H., & Koole, S.L. (2014) Internally directed cognition and mindfulness: an integrative perspective derived from predictive and reactive control systems theory. Front. Psychol., 5, 429.

Townshend, J.M. & Duka, T. (2001) Attentional bias associated with alcohol cues: differences between heavy and occasional social drinkers. Psychopharmacology, 157, 67–74.

Uddin, L.Q., Yeo, B.T.T., & Spreng, R.N. (2019) Towards a universal taxonomy of macro-scale functional human brain networks. Brain Topogr., 32, 926–942.

Verplaetse, T.L., Cosgrove, K.P., Tanabe, J., & McKee, S.A. (2021) Sex/gender differences in brain function and structure in alcohol use: A narrative review of neuroimaging findings over the last 10 years. J. Neurosci. Res., 99, 309–323.

Volkow, N.D., Wang, G.-J., Tomasi, D., & Baler, R.D. (2013) The addictive dimensionality of obesity. Biological Psychiatry, 73, 811–818.

Vollstädt-Klein, S., Loeber, S., Kirsch, M., Bach, P., Richter, A., Bühler, M., von der Goltz, C., Hermann, D., Mann, K., & Kiefer, F. (2011) Effects of cue-exposure treatment on neural cue reactivity in alcohol dependence: a randomized trial. Biol. Psychiatry, 69, 1060–1066.

Wechsler, D. (1997) WAIS-III, Wechsler Adult Intelligence Scale, Third Edition: WMS-III, Wechsler Memory Scale, Third Edition: Technical Manual. Psychological Corporation, San Antonio, TX.

Wiers, C.E., Stelzel, C., Park, S.Q., Gawron, C.K., Ludwig, V.U., Gutwinski, S., Heinz, A., Lindenmeyer, J., Wiers, R.W., Walter, H., & Bermpohl, F. (2014) Neural correlates of alcohol-approach bias in alcohol addiction: the spirit is willing but the flesh is weak for spirits. Neuropsychopharmacology, 39, 688–697.

Woo, C.-W., Krishnan, A., & Wager, T.D. (2014) Cluster-extent based thresholding in fMRI analyses: pitfalls and recommendations. Neuroimage, 91, 412–419.

World Health Organization (2019) Global Status Report on Alcohol and Health 2018. World Health Organization.

Wrase, J., Grüsser, S.M., Klein, S., Diener, C., Hermann, D., Flor, H., Mann, K., Braus, D.F., & Heinz, A. (2002) Development of alcohol-associated cues and cue-induced brain activation in alcoholics. Eur. Psychiatry, 17, 287–291.

Yeo, B.T.T., Krienen, F.M., Eickhoff, S.B., Yaakub, S.N., Fox, P.T., Buckner, R.L., Asplund, C.L., & Chee, M.W.L. (2016) Functional specialization and flexibility in human association cortex. Cereb. Cortex, 26, 465.

Zhang, R. & Volkow, N.D. (2019) Brain default-mode network dysfunction in addiction. Neuroimage, 200, 313–331.

Zhou, H.-X., Chen, X., Shen, Y.-Q., Li, L., Chen, N.-X., Zhu, Z.-C., Castellanos, F.X., & Yan, C.-G. (2020) Rumination and the default mode network: Meta-analysis of brain imaging studies and implications for depression. Neuroimage, 206, 116287.

